# An operational framework to map Essential Life Support Areas (ELSAs) for biodiversity, climate, and sustainable development

**DOI:** 10.1101/2024.11.25.625159

**Authors:** Oscar Venter, Jamison Ervin, Anne Lucy Stilger Virnig, Scott Atkinson, Marion Marigo, Di Zhang, Christina Supples, Enrique Paniagua, Lea Phillips, Richard Schuster, Xavier Llano, Genevieve Pence, Peter Arcese, Luizmar de Assis Barros, Aray Belgubaeva, Daniel Borja, Steve Brumby, Neil D. Burgess, Leticia Cardozo, Carlos Cordero Vega, Maria Veronica Cordova, Liliana Corzo, Esteban Delgado-Altamirano, Claudia Fonseca, Edward Game, Yvio Georges, Hedley Grantham, Daniel Guerra, Andrew Hansen, Greer Hawley, Naborey Hout, Berexford Jallah, KC Deepak, William Llactayo Leon, James Leslie, Dy Lihong, Casandra Llosa, Abu Rushed Jamil Mahmood, Tsepang Makholela, Mark Mathis, Cornelia Miller Granados, Jennifer McGowan, Rafael Monge, Nokutula Mhene, Violeta Muñoz-Fuentes, Sandra Neubert, Menaka Panta Neupane, Fabiola Nuñez Neyra, Diego Olarte, Emmanuel T. Olatunji, Dorine Jn Paul, Veronica Recondo, Gerty Pierre, Hugh Possingham, Susana Rodríguez Buriticá, Kanat Samarkhanov, Vijaya Singh, Arnout van Soesbergen, Talgat Taukenov, David Telcy, Cristina Telhado, E. Abraham T. Tumbey, Piero Visconti, Holger Zambrano, James Watson

## Abstract

Almost all countries are making increasingly bold commitments to halt and reverse biodiversity loss, minimise the impacts of climate change, and transition to more sustainable development. The effective achievement of many of these commitments relies on integrated spatial planning frameworks that are adaptable to national circumstances, priorities and capabilities. This need is formally recognized by Target 1 of the Kunming-Montreal Global Biodiversity Framework (GBF), which specifies that all areas should be under such planning. Here, we describe the development and application of an operational framework for national-level integrated spatial planning: Essential Life Support Areas (ELSAs). This framework facilitates the identification of areas that - if protected, restored, or sustainably managed - can support the achievement of national commitments to biodiversity, climate, and sustainable development. The process of mapping ELSAs relies heavily on leadership by national experts and stakeholders and the integration of spatial data using systematic conservation planning tools. We showcase the ELSA process carried out for Ecuador, where the use of real-time scenario analyses enabled diverse stakeholder groups to collaborate to assess national priorities for nature, climate, and sustainable development, view trade-offs and synergies, and arrive at a spatial plan to guide national action. ELSA presented an actionable approach for Ecuador, and 12 other pilot countries, to create a spatial plan aimed at fulfilling their national and international commitments to nature, including to the GBF.

## Introduction

Humanity is facing a global biodiversity crisis, widespread climate disasters, and numerous public health emergencies, all stemming from the degradation of nature by unsustainable land and sea use practices and the neglect of environmental responsibilities.^1^ At the same time, efforts are intensifying to reconcile planetary health with humanity’s burgeoning socio-economic demands. These conservation and sustainable development efforts are guided by cross-cutting international agreements, such as the Kunming-Montreal Global Biodiversity Framework (GBF), the Sustainable Development Goals (SDGs), and the Paris Agreement on climate. These agreements chart a course for planetary survival and represent a call to action endorsed by all their signatory Parties.

While environmental agreements are often first defined in international fora, they are implemented through regional, national and sub-national policy and legislation, which are actioned by designated institutions with the necessary resources and authority. ^2,3^ Some key national policies translating global commitments into national action plans include National Biodiversity Strategy Action Plans (NBSAP), national SDG targets and National Development Plans and National Adaptation Plans (NAP). Yet, in-depth reviews of national planning efforts reveal that spatial environmental data seldom feature in national plans.^2,4^ For example, in 2019 only 15% of NBSAPs included spatial information capable of guiding specific action on the conservation and restoration of biodiversity.^5^ Moreover, no available national climate strategy included an actionable map that could help guide where mitigation and adaptation measures could best harness nature-based climate solutions.^4,6,7^ This lack of actionable maps in national environmental plans is concerning, since both the benefits and costs of environmental actions are spatially heterogeneous and non-fungible, and not having guidance on where actions should take place will result in suboptimal outcomes.^6,8^

One possible reason for the sparse inclusion of actionable maps in past national biodiversity and climate plans is the lack of a generally accepted framework for undertaking integrated spatial planning with stakeholders to achieve commitments to nature, climate, and sustainable development. Spatial planning for biodiversity is well established in frameworks such as systematic conservation planning (SCP),^9^ which provides a structured decision support process to identify and implement conservation areas. Yet most applications of SCP and other frameworks tend to have at least one of three important limitations.

First, some approaches focus primarily on mapping only biodiversity values, e.g. Key Biodiversity Areas, biodiversity hotspots ^10,11^, without considering broader socio-environmental values such as ecosystem services, natural resource management, and sustainable development.^12,13^ Second, existing planning frameworks often view the world as binary, with areas set aside for protection or available for development ^14,15^. Yet, global commitments cannot be achieved through protected areas in isolation and, instead, they must be supported across land and seascapes through actions that, in addition to protection, restore and sustainably manage biodiversity.^16–18^ Most analyses showcase that the minimum area needed to secure biodiversity and ecosystem service outcomes are much larger than current protected area targets,^17,19,20^ so sustainable management and restoration efforts need to be embraced and planned alongside protected area efforts to secure biodiversity outcomes at scale. It is important that spatial planning frameworks explicitly prioritize these actions, and not solely identify places to protect.

Finally, and most critically, many existing conservation planning frameworks tend to adopt a set of international criteria,^11,21–23^ instead of adopting specific national criteria and land use constraints.^24,25^ Global planning and assessments are important for informing on how well humanity is progressing towards environmental commitments,^26^ identifying broadly important geographies ^27^ and informing resource allocation of internationally fungible conservation resources.^28^ However, these spatial planning methods are reliant on global data that may not be nationally validated or accepted, and which are often very coarse in resolution and suffer from sampling bias,^29,30^ requiring careful interpretation before being utilised in national and local planning.^31,32^ More importantly, the use of global spatial data and criteria assumes that internationally coordinated and standardized land use decisions are applicable at national scales, when conservation decision-making is complex and grounded in national and subnational planning contexts, policy and legal frameworks, and stakeholder needs.^33^ This has led to calls for more participatory planning that embraces local knowledge and regional contexts with those stakeholders that have both an intimate understanding of place and the agency to facilitate change.^34,35^

The need for participatory planning is clearly highlighted through Target 1 of the GBF. This target, agreed to by all 196 Parties, commits to “ensuring that all areas are under participatory and integrated biodiversity-inclusive spatial planning”.^36^ Such planning is of course not an end in and of itself, but rather it is a precursor to achieving the other commitments set forth in the GBF, and other environmental agendas. In particular, integrated spatial planning can support the identification of suitable locations for achieving targets related to area-based conservation (e.g., GBF Target 3) and ecosystem restoration (e.g., GBF Target 2).

Here, in support of national achievement of GBF Target 1 and other global commitments more broadly, we present an operational framework for integrated spatial planning that harnesses recent advances in SCP tools to identify what we term Essential Life Support Areas (ELSAs). ELSAs are areas that, if protected, sustainably managed, or restored, can support the achievement of existing national or subnational policy commitments to biodiversity, climate, and sustainable development. The ELSA framework was co-developed and stress-tested through partnerships between the United Nations Development Programme (UNDP) and governments in 13 pilot countries: Cambodia, Chile, Colombia, Costa Rica, Dominican Republic, Ecuador, Haiti, Kazakhstan, Liberia, Nepal, Peru, South Africa, and Uganda.^37^ After presenting the five steps of the ELSA framework and how they were applied in Ecuador, we conclude by discussing lessons learned during the stress testing across all 13 countries and outline how the ELSA framework can be utilised by nations to achieve current international and national commitments to nature, climate and sustainable development.

### Mapping Essential Life Support Areas in Ecuador

Ecuador was chosen as an ELSA pilot country for several reasons. First, it contains globally significant biodiversity and the second highest number of threatened species of any country.^38^ Second, it has a long history of efforts to achieve positive environmental outcomes. For example, Ecuador’s constitution of 2008 was the first in the world to recognize legally enforceable rights of nature.^39^ Third, the Ecuadorian government recognises the need to do more to secure its unique natural heritage. While Ecuador’s National System of Protected Areas (SNAP, by its acronym in Spanish) covers 26.2 million ha as of 2023, which is equivalent to 19.4% of its national territory, and the national incentive program of payments for biodiversity conservation (known as the SocioBosque Program in Spanish) conserves a further almost 1.6 million ha of private and community lands,^40^ there is a national recognition that many important areas for nature remain outside of the national conservation estate.^10,40,41^ National efforts to deliver on Ecuador’s commitments to the CBD are led by the Ministry of Environment, Water, and Ecological Transition (MAATE, by its acronym in Spanish), which works to integrate all national efforts to conserve at least 30% of land and waters across government, civil society, intergovernmental organizations, and Indigenous Peoples and local communities (IPLCs). Beyond protected areas, integrated spatial plans are needed to identify critical areas in Ecuador where sustainable management and restoration efforts can complement and connect protected lands.

Despite high technical capacity and strong political will, the country faces challenges to implement integrated spatial planning that are common to many developing, middle-income, and small nations. These include the lack of and uneven access to environmental spatial data;^29^ difficulty in formalizing the relationship between government and academia in support of environmental decision making, fragmentation of data across ministries, NGOs, and academic institutions; and challenges to integrated planning, implementation, and reporting across diverse sectors.^42^ To help address these challenges, in 2021, in partnership with the Ecuadorian government, UNDP convened partners in Ecuador to harness national expertise and spatial data to develop a national map of ELSAs to guide efforts to achieve key national commitments to nature, climate, and sustainable development.

ELSA’s were mapped through five distinct project steps (figure 1) which are founded on the theory of change for improved post-2020 (now known as the GBF) conservation outcomes proposed by Rice and colleagues.^43^ This provides a simple and logical sequencing of what needs to happen to achieve a conservation result^44^ and emphasizes the importance of producing a shared vision of desired results and actions at a national- and subnational-level, as well as the need to achieve a social and ecological balance. The steps are designed around a holistic, community-centered, context-specific, and adaptive approach to planning (Fig 1).

**Figure 1.**
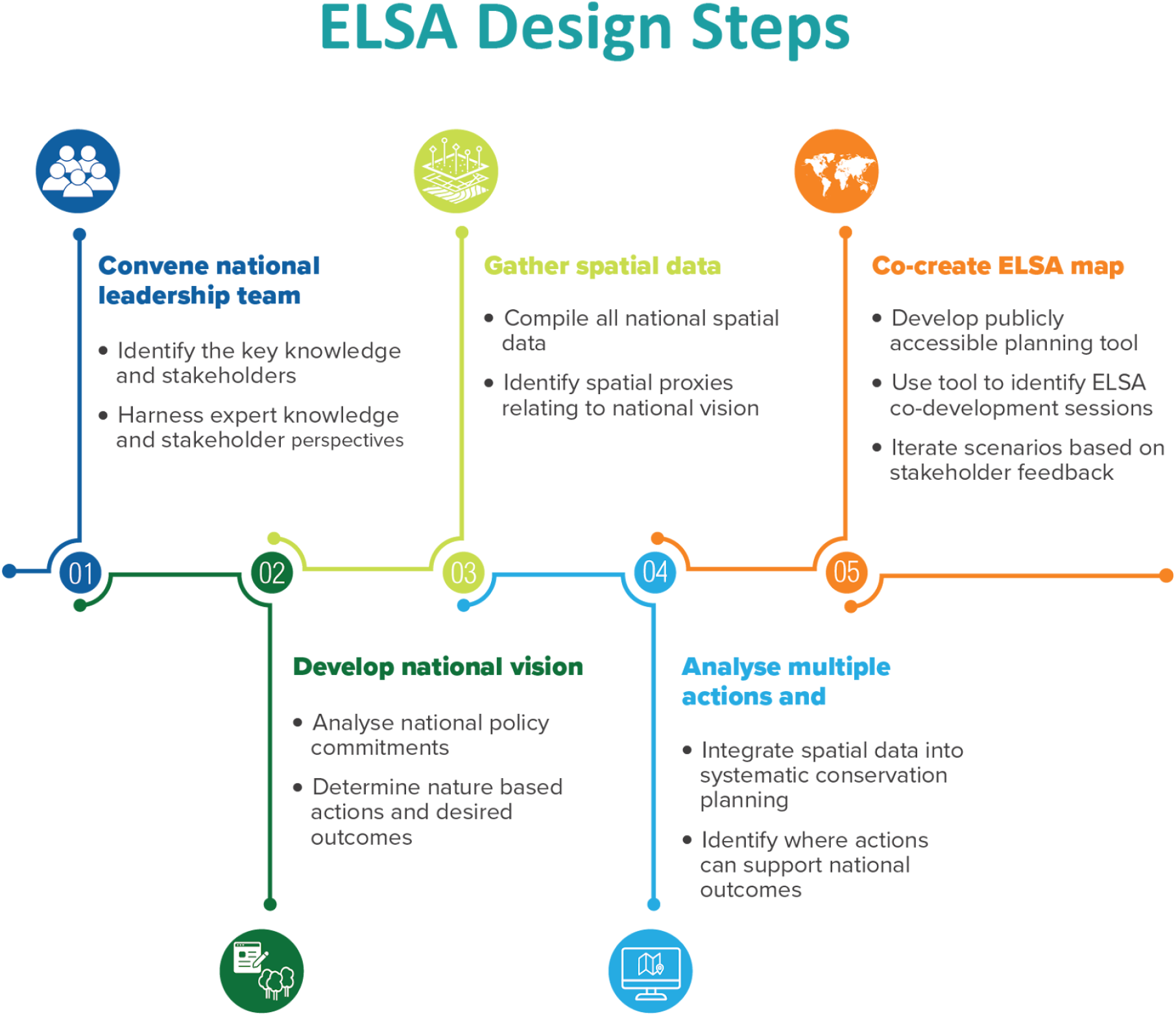
Five steps for identifying Essential Life Support Areas (Images adapted from Rice et al.^43^)

### Step 1: Convene national leadership team

The first step of the ELSA mapping process involves actively engaging experts with relevant knowledge and stakeholders with vested interest or influence in the outcome. Engaging these groups to become leaders in the co-design and application of the spatial planning process is essential, because it ensures that the resulting spatial plan is credible, trusted, and applicable in policy making. Broad participation also helps develop a community of practice around the common objective of data-driven environmental decision-making while nurturing champions to help integrate the outputs of this spatial planning process into national and subnational policy and action.

Regular and structured stakeholder engagement was undertaken via the creation of core and broader stakeholder working groups to ensure that the knowledge and values from national and local experts and stakeholders shaped the ELSA mapping process. In Ecuador, the UNDP Country Office acted as the convening partner to identify members of the core team for ELSA mapping. As the only environmental authority and sole water authority in Ecuador, MAATE was engaged to lead the core team. The core team, which included MAATE, the UNDP Country Office, and the UNDP Global Programme on Nature for Development, took primary responsibility for convening other stakeholders, acquiring national spatial data, making methodological decisions after consultation with stakeholders, and communicating the ELSA work more broadly.

The core team’s first task was to convene a broader stakeholder working group to provide a comprehensive perspective of national context and priorities to guide the ELSA mapping (Table 1). The stakeholder working group was made up of representatives from the Ministry of Agriculture and Livestock, Ministry of Tourism, Ministry of Foreign Affairs and Human Mobility, Ministry of Public Health, National Secretariat of Planning, National Risk and Emergency Management Service, Agency for Regulation and Control of Energy and Non-Renewable Natural Resources, Ministry of Economic and Social Inclusion, Military Geographic Institute, National Institute of Public Health Research, Geological and Energy Research Institute, National Institute of Meteorology and Hydrology, National Council for the Equality of Peoples and Nationalities, Foundation ALDEA, Foundation Charles Darwin, Pontifical Catholic University of Ecuador, and Ikiam Amazon Regional University. Indigenous Peoples and local communities were represented by the Confederation of Indigenous Nationalities of the Ecuadorian Amazon (CONFENIAE in Spanish), a regional indigenous organization that represents nearly 1,500 communities, and the Amazonian Network of Georeferenced Socio-environmental Information, a consortium of civil society organizations.

**Table 1.**
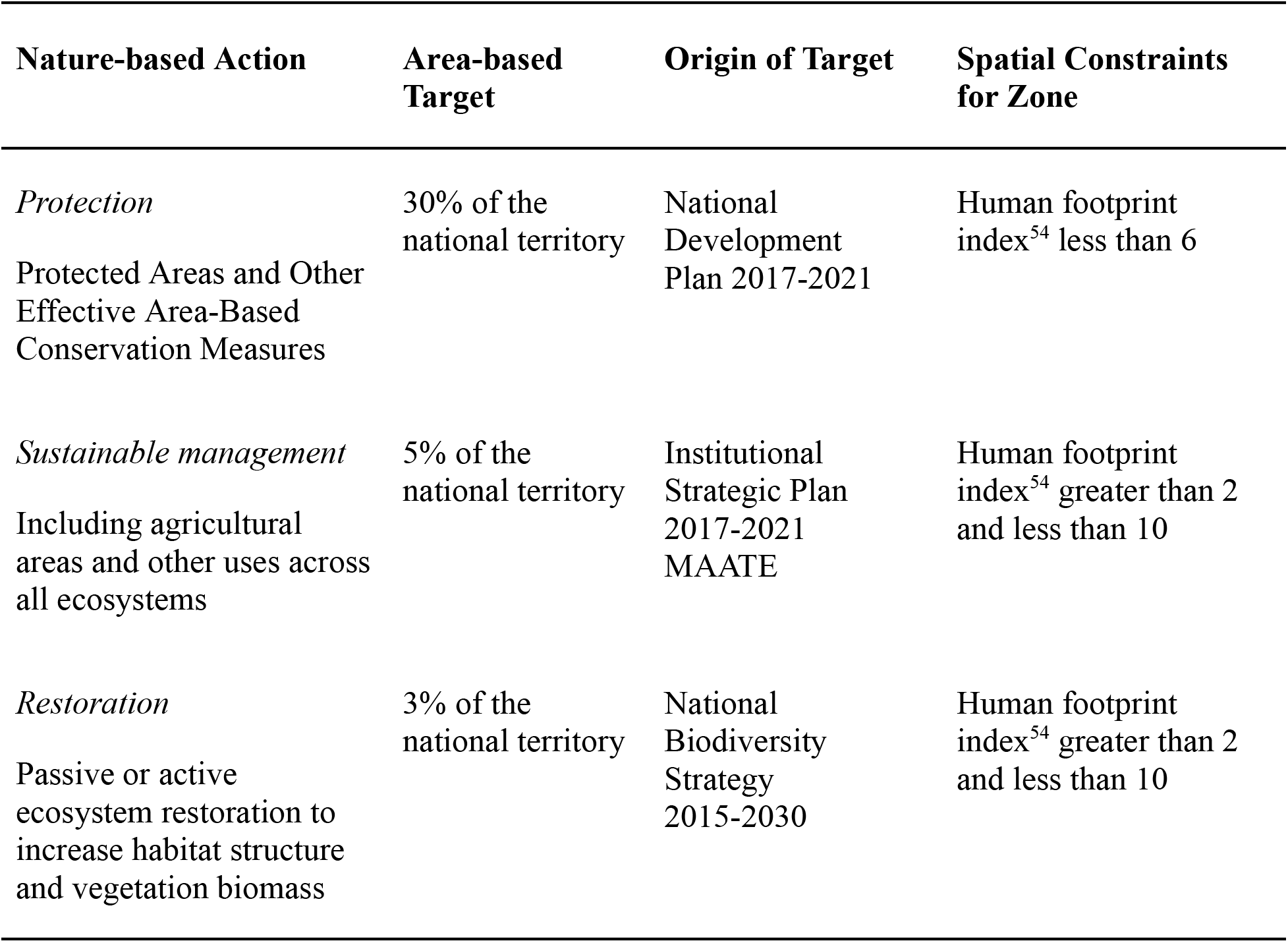
Nature-based actions and their area-based targets and spatial constraints (locations where each zone can be considered during the optimization).

| **Nature-based Action** | **Area-based Target** | **Origin of Target** | **Spatial Constraints for Zone** |
| --- | --- | --- | --- |
| *Protection*<br>Protected Areas and Other Effective Area-Based Conservation Measures | 30% of the national territory | National Development Plan 2017-2021 | Human footprint index<sup>54</sup> less than 6 |
| *Sustainable management*<br>Including agricultural areas and other uses across all ecosystems | 5% of the national territory | Institutional Strategic Plan 2017-2021<br>MAATE | Human footprint index<sup>54</sup> greater than 2 and less than 10 |
| *Restoration*<br>Passive or active ecosystem restoration to increase habitat structure and vegetation biomass | 3% of the national territory | National Biodiversity Strategy 2015-2030 | Human footprint index<sup>54</sup> greater than 2 and less than 10 |

The stakeholder group provided structured input through four stakeholder consultations, including a policy hackathon to support ELSA Step 2, a data hackathon to support Step 3, a methods review to support Step 4, and ELSA co-creation sessions to support Step 5, described in detail below. Across these four consultations and pre and post correspondence, key national stakeholders provided essential information to guide the design of Ecuador’s ELSA map. During the workshops, knowledge and values from the participants were harnessed via the IDEA protocole^45^ which minimizes ‘group think’, where individuals conform to the consensus view without engaging in critical discussion, and instead seeks to enable all individuals to provide input before aggregation of the expert and value-driven data.

### Step 2: Develop a national vision

The national vision for ELSA Ecuador was developed through a combination of a policy analysis and stakeholder engagement focusing on policy interpretation. The policy analysis relied on existing and validated national policy documents that the core team identified and collected. Examples of these policy documents included Ecuador’s National Development Plan, the National Water Plan, and the National Climate Change Strategy. These documents were then reviewed to extract all sections that detailed specific environmental commitments. Policy commitments were of two types. The first is a commitment to take a nature-based action, such as protecting or restoring a prescribed percentage of land area. The second is a specific commitment to achieve a conservation outcome, such as recovering an endangered ecosystem, ensuring clean water provision, or pursuing sustainable and climate-resilient agricultural practices.

Once the existing policy commitments had been compiled and reviewed, the first stakeholder consultation entailed a policy hackathon, which is an intensive collaborative exercise with the shared objective of solving policy challenges.^46^ During the policy hackathon, stakeholders were tasked with selecting the most critical policy commitments that described specific desired nature-based outcomes to form the national vision for ELSA in their country. To support this selection process, the compiled policy commitments were organized into three thematic areas -- biodiversity, climate change, and sustainable development -- enabling comparison of similar commitments and selecting the highest priority commitments representative of broad policy objectives. The prioritization of commitments across these three themes lays the foundation for discussions about the tradeoffs in planning for each independently versus in an integrated fashion during Steps 4 and 5 of the process. Through an iterative discussion and voting process, 10 priority policy commitments for inclusion in the ELSA mapping were agreed to (Figure 2).

**Figure 2.**
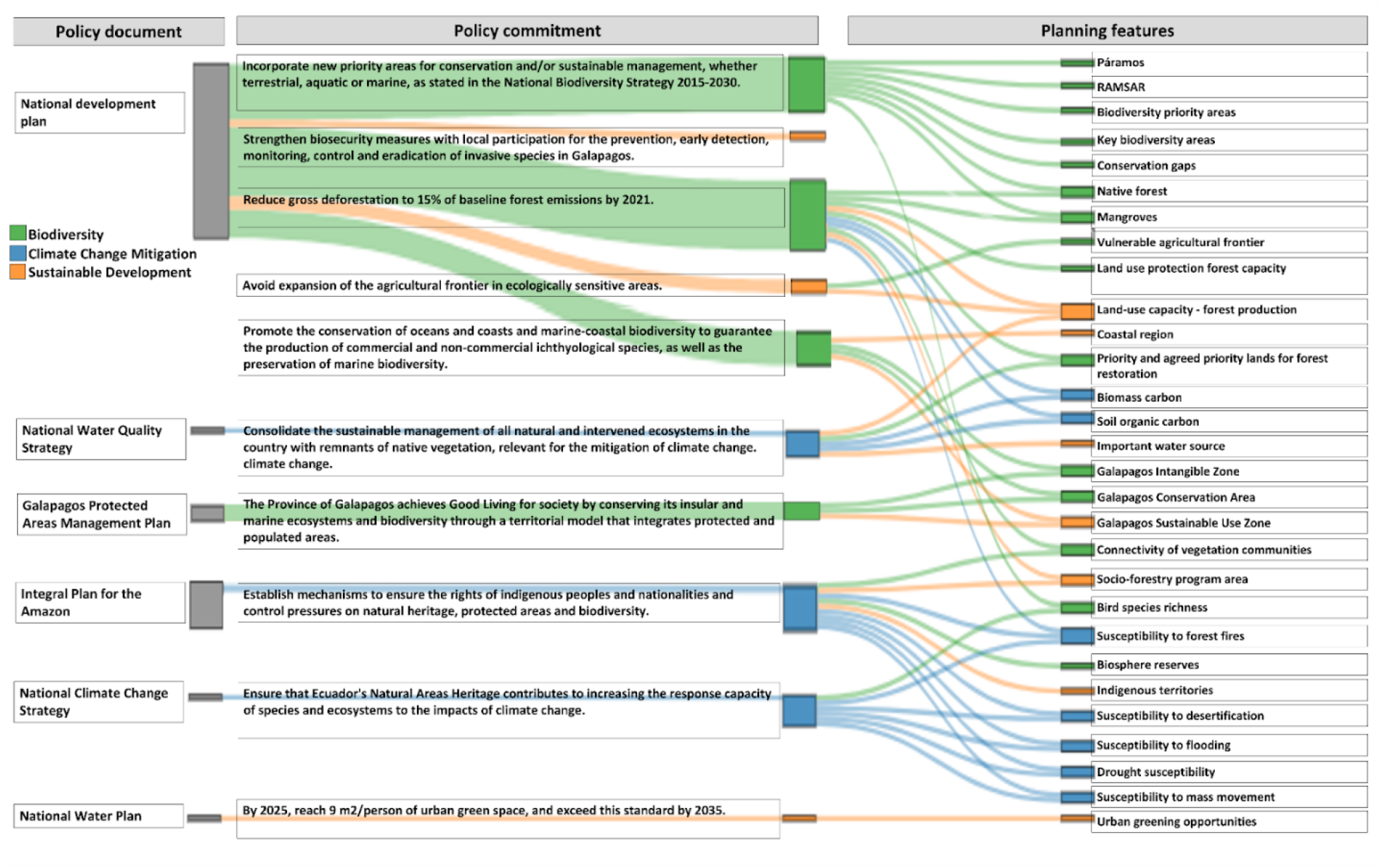
Stakeholders identified six national policy documents, from which they selected 10 priority policy commitments that defined the national vision for ELSA in Ecuador. They also identified national and global spatial data to serve as planning features and proxies for these commitments. Each of the spatial proxies were coded by stakeholders for biodiversity, climate mitigation and adaptation, or sustainable development for the purposes of evaluating tradeoffs among environmental outcomes (step 5).

Finally, stakeholders were tasked with reviewing the collected policy commitments and extracting any that set specific commitments for nature-based actions. Three commitments to nature-based actions emerged: 1) protection, which includes setting aside areas as nationally gazetted protected areas or other effective area-based conservation measures;^47,48^ 2) sustainable management, which involves reducing impacts of various land uses using various sustainable management techniques, such as reduced impact logging practices^49^ or permaculture techniques within agricultural landscapes;^50^ and 3) restoration, which involves restoring or rehabilitating degraded ecosystems by increasing the composition of native species or enhancing habitat structure and function.^51–53^ For each action, an area-based target was extracted directly from the policy commitments if it was given, or agreed to through unstructured group discussion (Table 1).

### Step 3: Gather spatial data

The third step is to identify spatial datasets that can serve as spatial proxies for the national vision and associated commitments. Environmental values, such as biodiversity and various ecosystem services, are not uniformly distributed across landscapes,^19,55,56^ and spatial data can identify where values are highest and overlapping.^23,57^ Datasets selected to map national commitments could include maps of biodiversity values, ecosystem services such as water security, food provision, and carbon, as well as datasets that map socio-economic values that relate to ecosystem functions such as agricultural suitability and industrial infrastructure. When identifying datasets, national data usually takes precedence over global data as it tends to better reflect national conditions, be viewed as more accurate by national users, and is more likely to be formally recognized for official use by governments.

Spatial data was compiled to meet two basic needs: 1) delineation of where nature-based actions, termed ‘zones’ in SCP, can occur and 2) spatial proxies for the policy commitments identified through the policy hackathon, termed ‘planning features’ in SCP. The identification of relevant national data was done through the data hackathon where national stakeholders and data experts identified existing national datasets on biodiversity, climate change, and sustainable development. Once the national datasets were identified, the core team engaged national data owners and relevant national institutions to secure the use of these data.

Once the national data collection was completed, all datasets were screened by the core team to ensure they were spatially explicit with area-based information, contained sufficient metadata, and were consistently mapped at the national level. When two or more datasets met these criteria and were thematically the same (e.g., two maps of forest cover), the national members of the core team identified which dataset should be retained, or if they should be combined using a consensus approach.^58^ Datasets were then further filtered to retain only datasets that could serve at least one of the two data needs, which were mapping possible locations for nature-based actions or serving as direct proxies for each of the 10 priority policy commitments. Finally, global datasets were extracted from the UN Biodiversity Lab (www.unbiodiversitylab.org) inventory for any remaining data needs not covered by national data, and reviewed with the core team to ensure they were nationally acceptable.

A total of 28 national spatial datasets and three global datasets were selected for use in mapping ELSAs in Ecuador (Fig 2, Table S1, Figures S1-S29). The primary dataset used to map the potential location of each nature-based action was national data on human footprint ^54,57^ (Table S1). Datasets to map planning features spanned coarse filter proxies for biodiversity, such as nationally identified important ecosystems (e.g., paramos, mangroves, Ramsar sites) and fine filter maps (e.g., Key Biodiversity Areas and Biodiversity Priority Areas). Other datasets represented important spatial proxies for opportunities to either mitigate or adapt to climate change (e.g., soil organic carbon, drought risk), while others represented ecosystem services important for sustainable development (e.g., land use capacity, water sources, indigenous territories). To evaluate trade-offs among broad conservation goals, each dataset was identified as supporting particular policy commitments, as well as one of biodiversity, climate change, or sustainable development (Figure 2).

### Step 4: Analyse multiple actions and outcomes

The fourth step is to use systematic conservation planning to analyse land use zones and outcomes. SCP is used to optimize spatially-explicit conservation actions to promote the persistence of biodiversity and other natural features in situ.^9,20,59^ SCP involves a transparent and objective process of setting clear goals and objectives, and subsequent planning for conservation actions that meet them.^60^ SCP was originally developed to identify alternative proposed networks of protected areas. More recently it has evolved to consider multiple nature-based actions and objectives beyond biodiversity, making it suited for engaging with the complexity of integrated spatial planning across landscapes and nations. ^61–63^ SCP was used to analyse all nature-based actions and planning features at once, thus capitalizing on spatial synergies across all policy commitments when identifying ELSAs. In addition to integrating multiple commitments, SCP enables diverse stakeholder groups to weigh the relative importance of the various planning features, view trade-offs that result from conflicting priorities, and foster dialogue around cross-sectoral collaboration and implementation.

The ELSA analysis uses the prioritizr software library to run the SCP analyses. ^64,65^ The prioritizr package is conceptually similar to the widely used planning software Marxan,^66^ but differs in its implementation of integer linear programming techniques^67^ instead of simulated annealing as the solving algorithm. The linear programming approach can solve large problems (>1 million planning units) faster than other approaches, allowing real-time analysis with stakeholders. Moreover, it supports a broad range of objectives, constraints, and penalties that can be used to customize conservation planning problems to the specific needs of a conservation planning exercise.

For each zone, its contribution to representing each planning feature is set using a zone contribution score (Table S2). For example, if a feature falls 100% inside the ‘Manage’ zone and the zone contribution for that feature is ‘0.5’, the feature representation score would be 50% of its original mapped extent. On the other hand, if a feature falls completely within the ‘Protect’ zone and the area has a ‘1.0’ zone contribution for the feature, its representation score would be 100%. Also, for each zone, a map is used to identify where it can be considered within the optimization, based on national input (Table S3, Fig S30).

The maximum utility optimization function (https://prioritizr.net/reference/add_max_utility_objective.html) within prioritzr is used for its ability to find locations for the nature-based actions that maximize the total representation of planning features, accounting for zone contributions, with the relative importance of each planning feature controlled through a weighting parameter. The maximum utility function differs from the more widely used minimum set functions, where specific representation targets are set for each planning feature (e.g., protect 25% of a species range). The maximum utility function was used for two reasons. First, the diversity of planning features (e.g., important water source areas, and connectivity of vegetation communities) presented challenges to setting evidence-based targets for all features. Second, Ecuador, like many countries, has clear targets for zones (e.g., protecting 30% of land areas), which is a core parameter for the maximum utility function.

To promote equity in representation across planning features, we conduct a pre-calibration process in which a script: 1) weights all planning features equally, evaluating how well each feature is represented in the solution (e.g., its maximum utility); 2) weights each feature as 1 while setting all other features to 0, and again solving the problem to see the impact of that feature’s weight on the overall solution (e.g., its maximum representation); and 3) finally, enters a calibration loop where it iteratively adjusts the weights based on the difference between the maximum utility and maximum representation for each feature, aiming to minimize the difference (delta) between these values and leading to a more equitable representation across all features. These pre-calibration weights then serve as our starting weights on the server backend for the ELSA co-creation sessions (step 5).

### Step 5: Co-create the ELSA map

The final step is to use the SCP tool to co-create the ELSA mapping analysis through real-time iterative scenario analyses with stakeholders,^63^ which is a departure from the model of global experts sharing the results with stakeholders once they are created. As the ELSA process integrates multiple, often competing, priorities in a given country, leadership from national experts and stakeholders is key for evaluating trade-offs across scenarios and iterating maps to identify a final product that best meets the diverse objectives of the national vision.

To allow full involvement of the core team and broader stakeholder group within the ELSA mapping process, and avoid the common scenario where a group of technical experts independently run the SCP optimizations, we developed a simple online user interface for the prioritizr optimization package preloaded with Ecuador’s spatial data (http://elsa.unbiodiversitylab.org/Ecuador_ELSA/). The tool allows data visualization, setting of targets and weights, real-time (∼2 minutes) optimization runs, display of the resulting ELSA action maps, and tabular analysis of the results. The co-creation of the final ELSA map was done using this tool through two sessions involving all participants from the core team and broader stakeholder group.

In the first session, weights for each planning feature were assigned by the broader stakeholder working group and non-UNDP members of the core group. During this weighting session, each spatial dataset was shown to participants, and its source, characteristics, and meaning were discussed. Stakeholders were then asked to give the dataset a weight between zero and ten quantifying their perspective of how important the planning feature should be in guiding the identification of ELSAs and supporting national environmental commitments. The average weight across the stakeholders was then used to create the initial ELSA map in the second co-creation session.

The initial ELSA action map serves to identify areas for each action (protection, sustainable management, restoration) to achieve the area-based targets (Table 1) in a way that maximizes the representation of all planning features, given their weights. To evaluate the trade-offs of integrated planning for this first map, separate ‘siloed’ planning runs are performed independently focusing only on planning features identified as 1) biodiversity, 2) climate, and 3) sustainable development (Figure 2). The representation of each planning feature in the initial ELSA map was then measured against its representation within the siloed maps. All planning features with a drop in representation of 15% or greater in the ELSA integrated planning map were then flagged and this ‘trade-off’ of integrated planning was discussed as a group.

A voting exercise was undertaken to determine if the weights should be further adjusted to increase the representation of each or certain planning features that experienced this drop in representation. For four out of the five planning features experiencing a greater than 15% drop in representation, the majority of participants voted to increase their weights, and for these, the weights were then increased by 50% (predetermined as a level that led to an observed increase in representation) and a new ELSA action map was produced. This process was repeated until voting indicated no further changes to the weights.

This agreed ELSA map (Figure 3) shows where actions can most effectively achieve the greatest impact across all planning features while minimizing unacceptable tradeoffs of integrated spatial planning.^68^ This map of ELSAs outlines an ambitious expansion of protected areas and other effective area-based conservation measures from 25.5% of the land area covered by existing protected areas to 30%. It also outlines critical areas to pursue sustainable management practices (5% of land areas) and ecosystem restoration (3% of land areas) to achieve multiple environmental, climate, and sustainable development outcomes.

**Figure 3.**
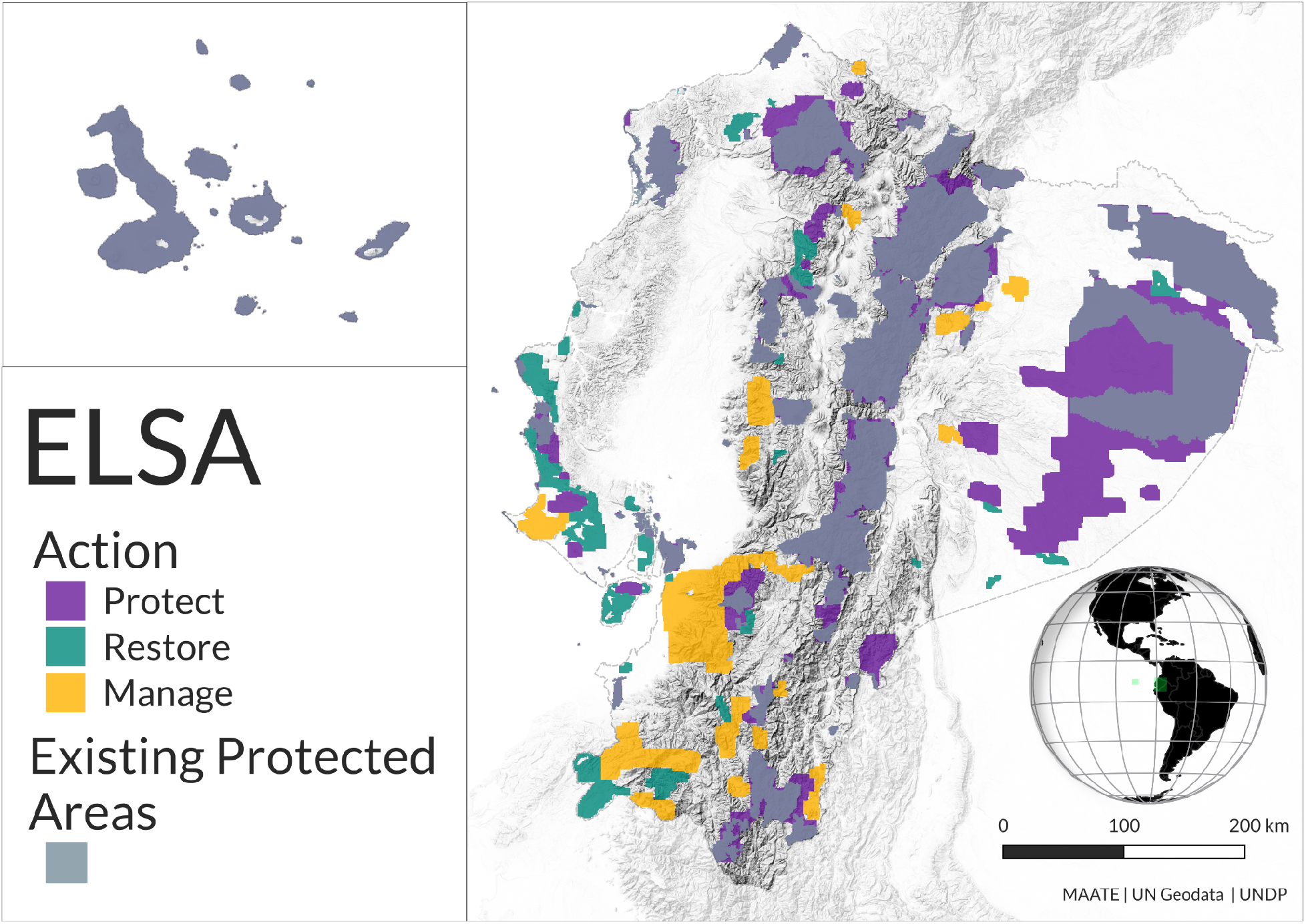
Adopted ELSA map identifying where achieving 30% protection, 5% management and 3% restoration can maximize combined representation across all planning features and achieve Ecuador’s national vision for nature.

An important component of the ELSA map is the contribution across zones in representing planning features (Figure 4). Some planning features are only represented within a single zone. For instance, areas susceptible to desertification are represented only within the ‘protect’ zone, whereas important water sources are only represented within the sustainable management zone. However, the majority of planning features are represented across all zones (Figure 4), highlighting the importance of considering a range of zones for achieving the diversity of national commitments to nature, climate, and sustainable development.

**Figure 4.**
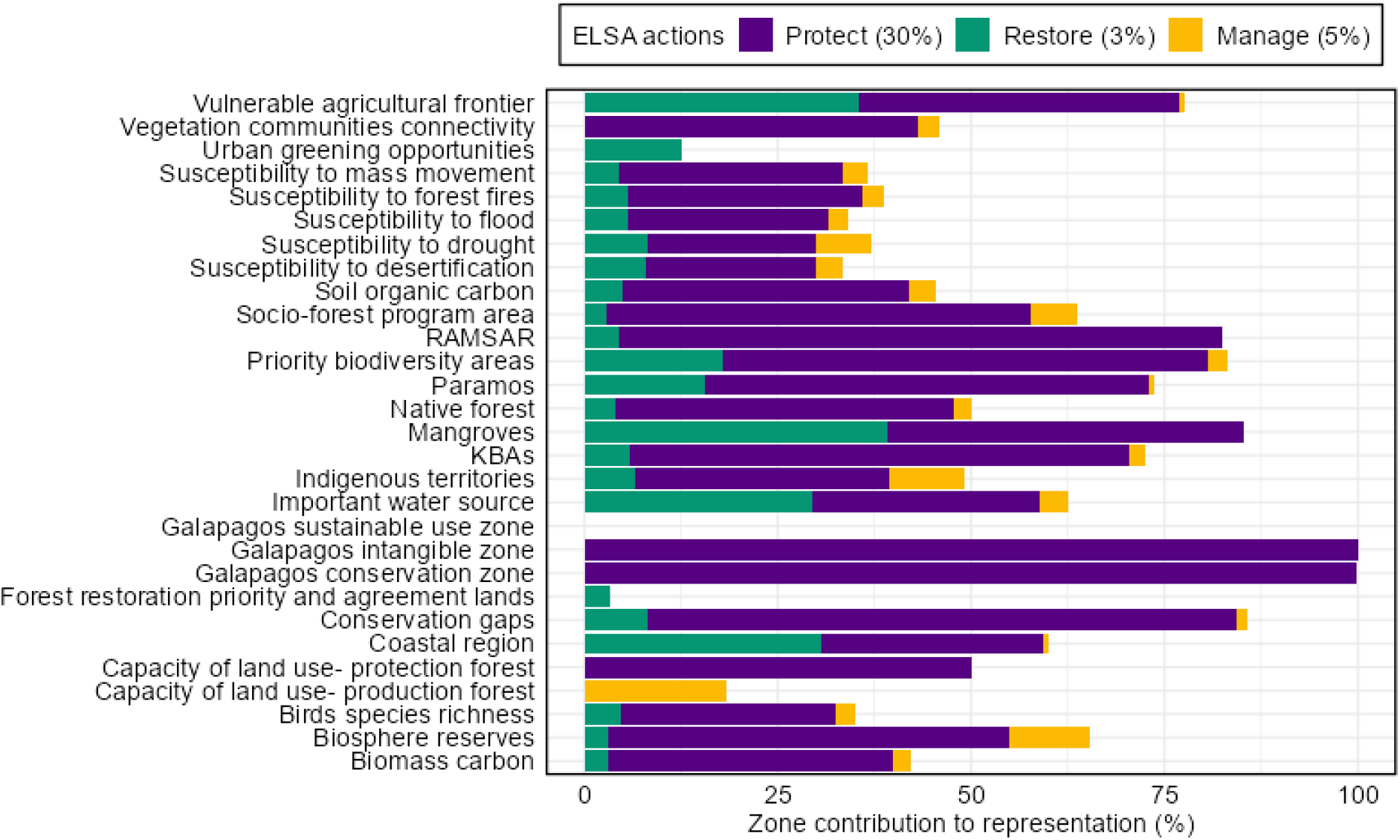
The contribution of each zone to the representation of planning features in the ELSA map. Representation (%) of a planning feature is the sum of its coverage per zone (‘Protection’, ‘Management’, and ‘Restoration’) multiplied by each feature-specific zone contribution (Table S2), relative to the total possible amount of the feature across the whole planning area.

## Discussion

The impetus behind ELSA’s development was the observed lack of actionable maps within national policy documents,^4–6^ as well as the growing call for the adoption of nationally relevant integrated spatial planning,^36^ such as through GBF Target 1. The ELSA framework was built through guidance and input from national experts and stakeholders as critical leaders and beneficiaries of integrated spatial planning.^17,69^ It also recognises that while the actual transformation in land use and sea use activities will likely occur at more localised scales,^15^ it is primarily through national-scale planning that the necessary governance arrangements can be put in place to operationalize environmental agendas, assess trade-offs, and ensure that citizens and wider society, including the private sector and Indigenous Peoples (IPs) and local communities (LCs), can actively participate.^70^

In Ecuador, the ELSA mapping project convened participants from 18 government ministries and national institutions, as well as Indigenous Peoples and civil society organizations. These participants led the defining of the national vision for environmental outcomes through a review and prioritization of existing national environmental commitments. It is interesting to note that though policy commitments were extracted from Ecuador’s NBSAP, none of these were identified by participants as the most critical for identifying ELSAs. Instead, five priority policies were drawn from the National Development Plan, though all of these described biodiversity or other environment-related aspects of development, while others were drawn from plans focusing on critical ecosystems and services (e.g., Water Quality Plan). Yet, when participants weighted the actual individual spatial data layers (planning features), they on average gave the highest weights to layers representing biodiversity values. This ability to recognize environmental outcomes across broad environmental agendas shows that diverse groups of national stakeholders can come together and generate maps to prioritise the protection of the environment broadly. The resulting agreed ELSA map represents an ambitious vision developed by an inclusive group of stakeholders and experts, and has the potential to inform ongoing environmental action in the country. Led by MAATE, the ELSA map is currently being updated to contribute to the country’s NBSAP to guide Ecuador’s action on targets under the GBF.

Between 2020 and 2023, ELSA mapping was undertaken in an additional 12 pilot countries where it proved to be flexible and adaptable to national contexts.^37^ Like Ecuador,^71^ the primary contribution of the ELSA projects was the integration across biodiversity, climate, and sustainable development agendas and the consideration of actions in landscapes beyond protected areas, as well as an approach that fully harnesses national policy vision and national expertise and values. However, the national vision defined in each case presented a unique set of values and challenges. In South Africa,^72^ a country that has pioneered spatial planning science and applications over the past 30 years,^21^ the ELSA methodology provided a novel extension to enable the creation of a multi-zone analysis that directly reflected national nuances in policy approaches. Uniquely, in South Africa the ELSA methodology included zones for 1) protection, 2) avoiding loss, 3) reducing pressures, 4) restoring, and 5) fostering urban adaptation, which reflected the policy and legal instruments for environmental outcomes and land management in the country. In Costa Rica, the ELSA project was leveraged as a key component in the development of the National Climate Adaptation Plan 2022-202,^73^ contributing a strong focus on spatial data to map climate vulnerabilities and facilitate climate change adaptation efforts, while providing important co-benefits for biodiversity and sustainable development. In Colombia, the ELSA methodology was applied at a subnational scale for the first time, addressing the issue of water security alongside concerns for biodiversity and climate change adaptation within the Central Region of Colombia, home to the capital city of Bogota, and nearly 15 million people. The ELSA methodology and the resulting map demonstrated to national and regional policymakers the critical role of the páramos areas for water provision to densely populated cities and became a critical part of the Water Security Plan for the Central Region.^74^

One of the deliberate principles in the ELSA framework is that it needs to be nationally derived and not determined solely by international criteria; using existing national policy to structure ELSA serves to enhance the national support for the resulting outputs. Existing national policy documents represent a nationally-validated vision for environmental outcomes for shaping ELSA analyses. As more countries generate ELSA maps, it will be important to determine if cumulatively these national maps sufficiently support global commitments set in biodiversity, climate, and sustainable development agendas. This could be done through global stocktakes conducted by the CBD (e.g., Global Biodiversity Outlooks) or similar undertakings for the UN Framework Convention on Climate Change.

The pilot testing of the ELSA framework consistently demonstrated the importance of balancing the complexity of spatial analysis with the need for broad multi-stakeholder engagement, whilst still providing sufficient detail for real-world context and implementation.^75^ A key tool to allow this was the simplified user interface that was developed for each country to run optimizations and interpret ELSA maps in real-time through live co-creation sessions. The tool allowed all data inputs to be viewed by participants, as well as for the resulting ELSA maps to be panned and zoomed to focal areas that participants knew well, serving as something of a preliminary ground-truthing of results. However, the software development and data preprocessing necessary to establish the user interface tools have required the UN Nature for Development team to remain engaged with the technical aspects of subsequent national updates to ELSA analyses.

A key question for planning efforts that include collaboration across international organizations and national stakeholders is the level of national ownership (Table 2). Ultimately, the ELSA process aspires to enable national experts and stakeholders to independently conduct new and updated national ELSA maps. This would include national experts convening the relevant parties to define the national vision, selecting and processing input data, developing the SCP analysis and planning software, and using it in stakeholder workshops to evaluate planning scenarios.

**Table 2.**
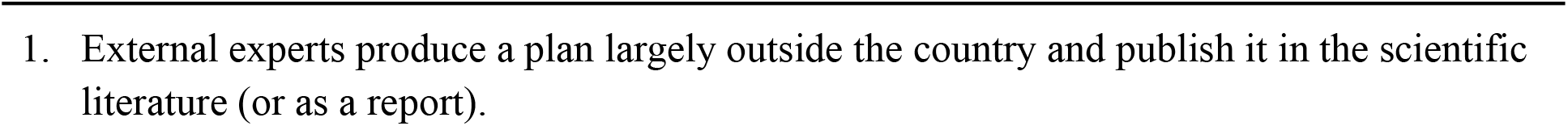
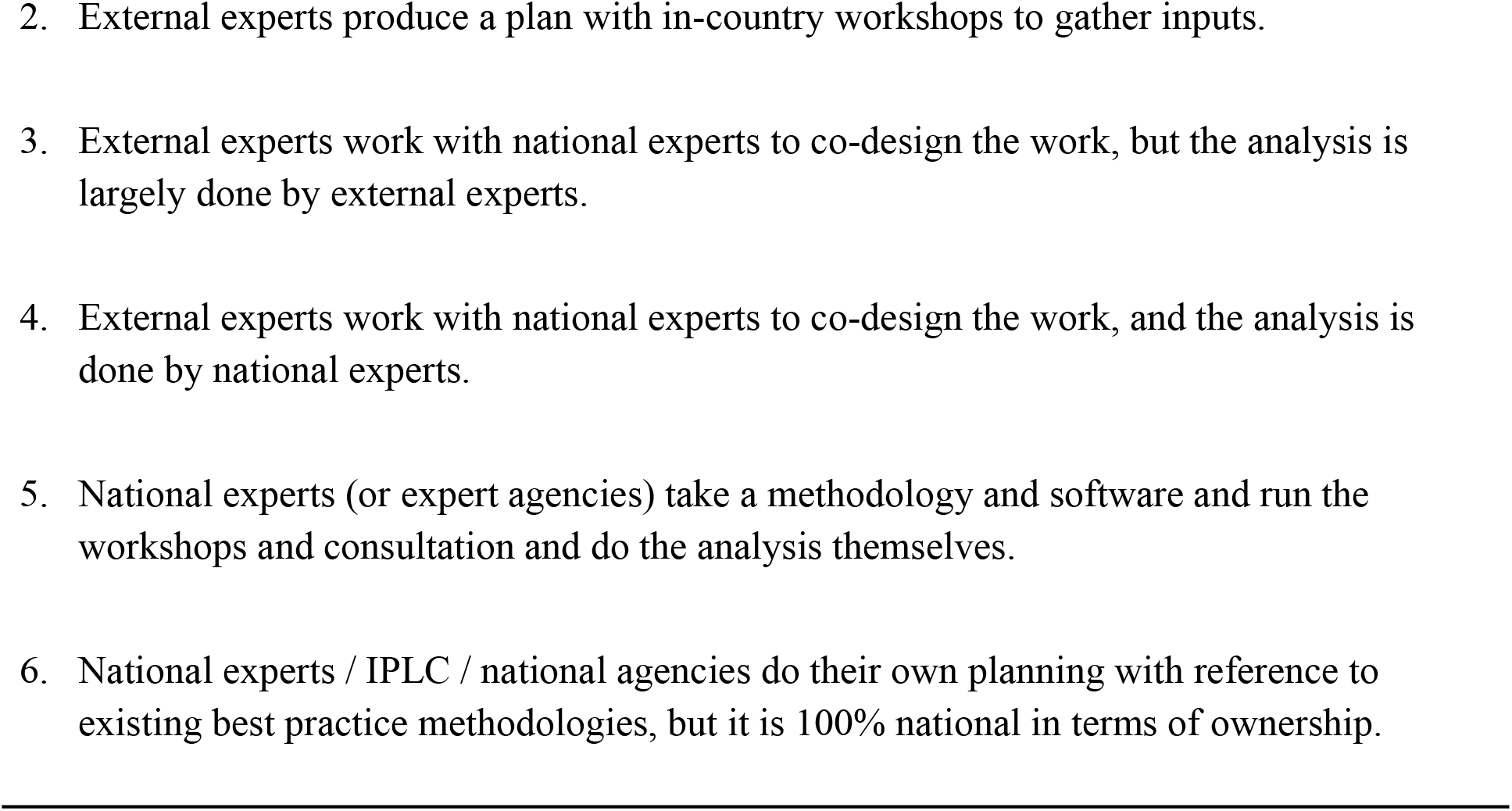
General framework to describe increasing levels of national ownership over a spatial planning exercise. To date, ELSA mapping has typically consisted of level 3 and level 4 national ownership.

This would be viewed as the highest level of national ownership (levels 5 and 6 in Table 2). To date, ELSA projects have achieved a level 3 and in some cases a level 4 of national ownership, with national experts and stakeholders leading most design decisions, but the UNDP Nature for Development team performing the majority of the technical analyses. This presents limitations for both national ownership of the results and the ability of national experts to undertake routine planning updates.

Thus far the ELSA team has worked around the technical barriers by continuing to support national ELSA updates. For instance, the effort currently underway in Ecuador to update their ELSA map to better support the new commitments they have made under the Global Biodiversity Framework in their updated NBSAP is supported by the UNDP Nature for Development team. However, scaling of the ELSA framework to all interested parties will require the development of technically simpler and more accessible tools. To do this, a new tool for rapid ELSA analyses using global data is under development by the UNDP Nature for Development team and United Nations Environment Programme World Conservation Monitoring Centre (UNEP-WCMC). Future applications of the ELSA framework are also striving to involve more diverse stakeholders, most notably the private sector and indigenous peoples and local communities, which play a large role in environmental outcomes.^76^ Future developments would also do well to include more land use activities, including mining and energy development and associated infrastructure networks, as it is increasingly recognised that many of these human activities need to be better planned.^77^ Finally, the ELSA framework should be equally applicable in the marine realm, where multiple stakeholders, values and sea-use zones all depend on sound planning,^78,79^ and efforts are under way to identify a case study to apply ELSA to the marine realm.

## Conclusion

The planning and decisions taken this decade will determine if our diverse planetary crises will overwhelm our ability and the ability of ecological systems to adapt, or if they will be addressed through effective and informed action. Biodiversity-inclusive spatial planning, such as implemented through the ELSA approach, has the potential to be one of several tools to support action during this critical window of time.

Our collective hope is that the centrality of GBF Target 1 will help drive the generation of more national participatory biodiversity-inclusive spatial planning processes. However, it is widely acknowledged that there is a gap in available methods, data, and a community of practice around this type of spatial planning to support Parties to easily take action.^80^ The ELSA approach offers one way to do this through a formalized process for fostering cooperation around spatial planning. It has the potential to support Parties to achieve not only GBF Target 1 (integrated spatial planning), but also to create maps that show where implementation of the area-based GBF Targets 2 and 3 (ecosystem restoration and conservation) can provide the best co-benefits for related biodiversity, climate, and sustainable development commitments, including GBF Targets 4-12, which are all mappable and nature-based. Innovations in spatial planning such as ELSA can help Parties to take more effective action to transform society’s relationship with nature.

## Acknowledgements

This work was supported through funding from the Global Environmental Facility (GEF), Gordon and Betty Moore Foundation, One Earth, Sustainable Markets Foundation, and the Swedish International Development Cooperation Agency (Sida). In addition to our co-authors, we would like to acknowledge the input and advice from Kulee Keculah (Liberia), José Luis Naula (Ecuador), and Andrew Skowno (South Africa), as part of our national level implementation of the ELSA approach, and Justina Ray as part of our Expert Advisory Committee. We would also like to acknowledge the UN Biodiversity Lab, from which we identified global data that could be used to fill national data gaps, when validated by national experts.

## Data availability

*The datasets used to obtain the ELSA map of Ecuador are available at https://github.com/ELSA-UNDP/elsa_national_public. The source and access of the original national data from Ecuador are provided in the Supplementary Materials*.

## Code availability

*We provide a publicly available sample code repository for an example ELSA Rshiny application at https://github.com/ELSA-UNDP/elsa_national_public, providing both data and instructions on tool set-up, allowing users to run the application on their local computer using free and open-source solvers in the R language*.

## Supplemental information

Document S1. Figures S1–S30 and Table S1-S3

## Notes

### Competing Interest Statement

The authors have declared no competing interest.

