## Supplementary materials for "An operational framework to map Essential Life Support Areas (ELSAs) for biodiversity, climate, and sustainable development"

Table S1 Input data used in ELSA Ecuador

| Group | Theme | Name | Data Source | National/ Global |
|---|---|---|---|---|
| Planning features | Biodiversity | Native Forest | Ministry of Environment (MAE), 2018 | National |
| | | Mangroves | MAE, 2017; Charles Darwin Foundation (FCD), 2015 | National |
| | | Paramos | MAE, 2018 | National |
| | | RAMSAR Sites | MAE, 2017 | National |
| | | Forest restoration priority and agreement lands | MAE, 2014; 2019 | National |
| | | Forest for protection | MAG, 2019 | National |
| | | Priority biodiversity areas | CONDESAN/MAE, 2013 | National |
| | | KBAs | Birdlife, 2020 | Global |
| | | Conservation gaps | CONDESAN/MAE, 2013 | National |
| | | Biosphere reserves | MAE, 2018 | National |
| | | Bird species richness | MAE, 2013 | National |

| | | | |
|---|---|---|---|
| | Connectivity of vegetation communities | MAE, 2012 | National |
| | Vulnerable agricultural frontier | CONDESAN/MAATE, 2013; MAG, 2019; MAE, 2017; MAE, 2018; Birdlife, 2020 | National |
| | Galapagos intangible zone | FCD, 2017 | National |
| | Galapagos conservation zone | FCD, 2017 | National |
| Climate Change Mitigation | Biomass carbon | Spawn et al., 2020 | Global |
| | Soil organic carbon | Ministry of Agriculture and Livestock (MAG), 2017 | National |
| | Susceptibility to desertification | Military Geographic Institute (IGM), 2013 | National |
| | Susceptibility to flood | IGM, 2015 | National |
| | Susceptibility to forest fires | National Service of Risk Management and Emergencies (SNGRE), 2015 | National |
| | Susceptibility to drought | IGM, 2015 | National |
| | Susceptibility to mass movement | SNGRE, 2011 | National |
| Sustainable Development | Urban-greening opportunities | National Institute of Statistics and Census (INEC), 2014 | National |

| | | | | |
|---|---|---|---|---|
| | | Socio-forest program area | Ministry of Environment, Water and Ecological Transition (MAATE), 2020 | National |
| | | Forest for production | MAG, 2019 | National |
| | | Indigenous territories | Socio-environmental Amazon (AMAZONIA), 2020 | National |
| | | Important water source | MAATE, 2020 | National |
| | | Coastal region | Oceanographic and Antarctic Institute of the Navy (INOCAR), 2017 | National |
| | | Galapagos sustainable use zone | FCD, 2017 | National |
| Lock-in | Protected areas | Protected areas of Ecuador | MAATE, 2020 | National |
| Zones | Input for zones | Human footprint index | Aragon et al., 2021 | National |

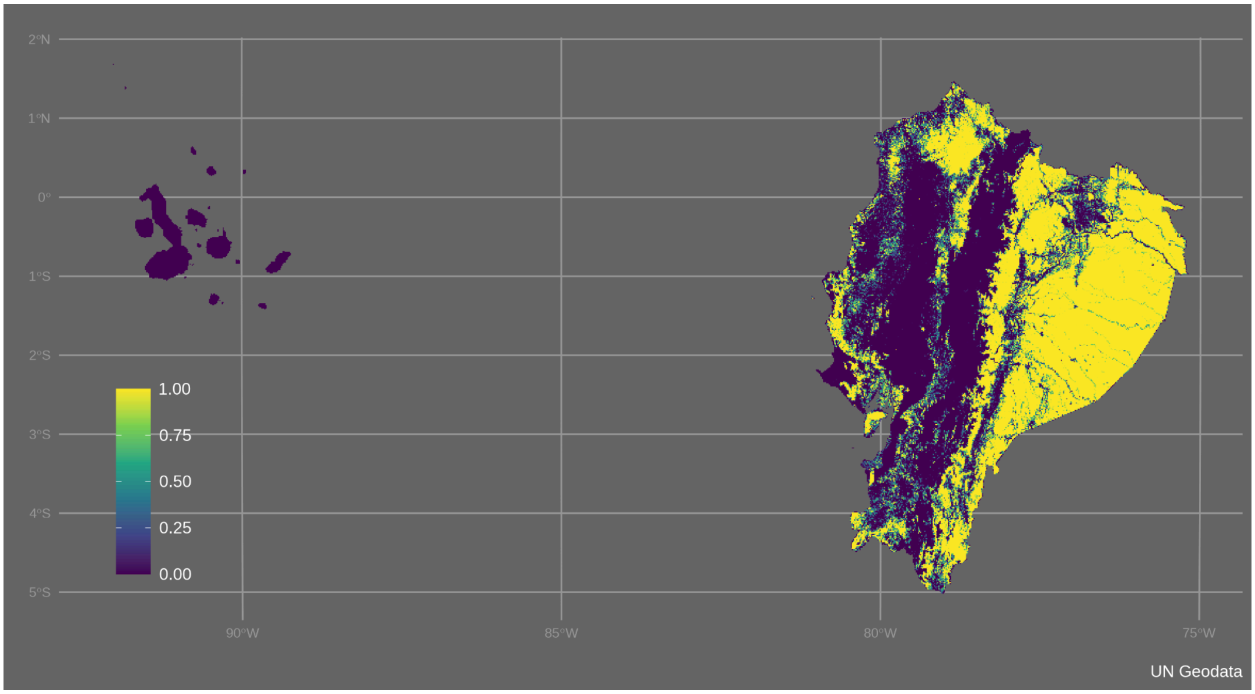


Figure S1. Planning features of ELSA Ecuador - native forest

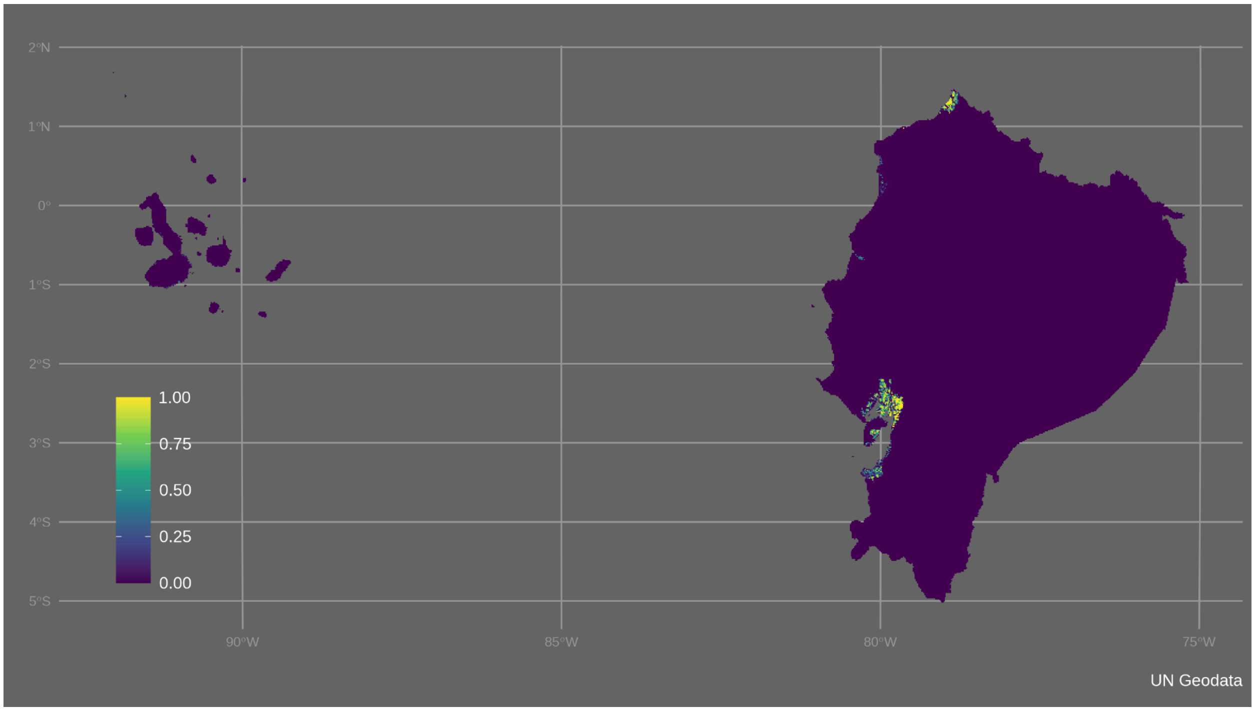


Figure S2. Planning features of ELSA Ecuador - mangroves

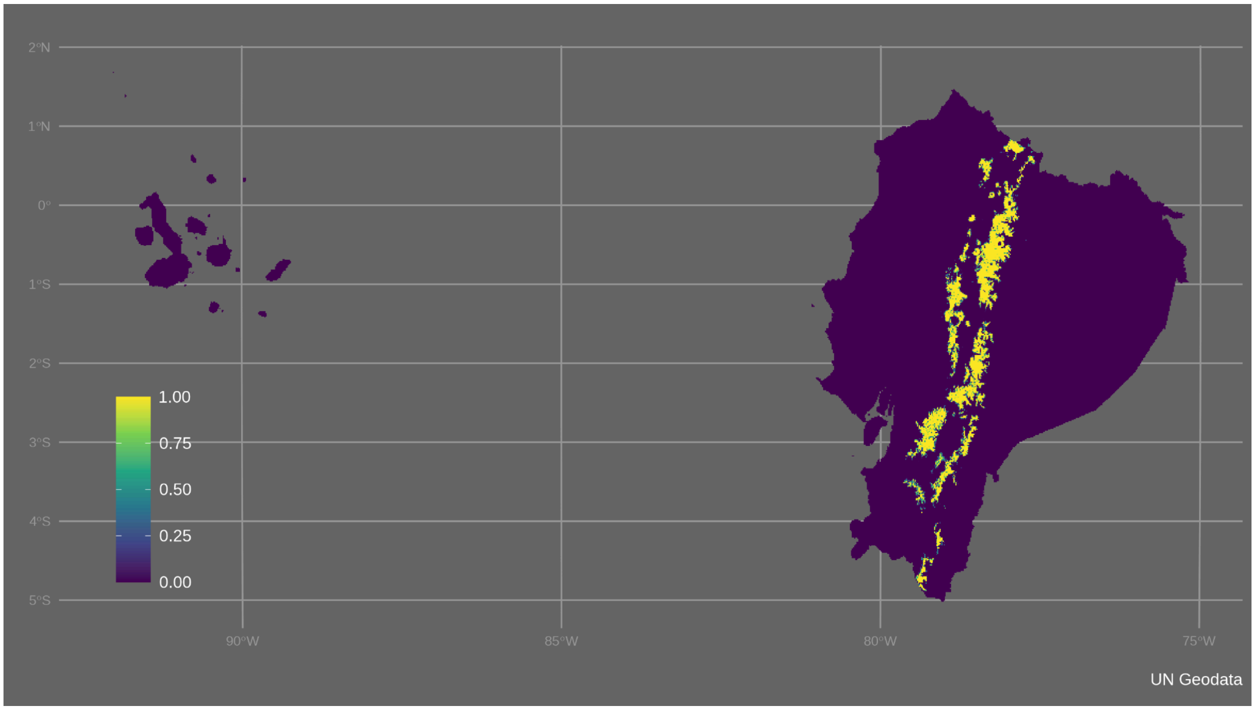

Figure S3. Planning features of ELSA Ecuador - paramos

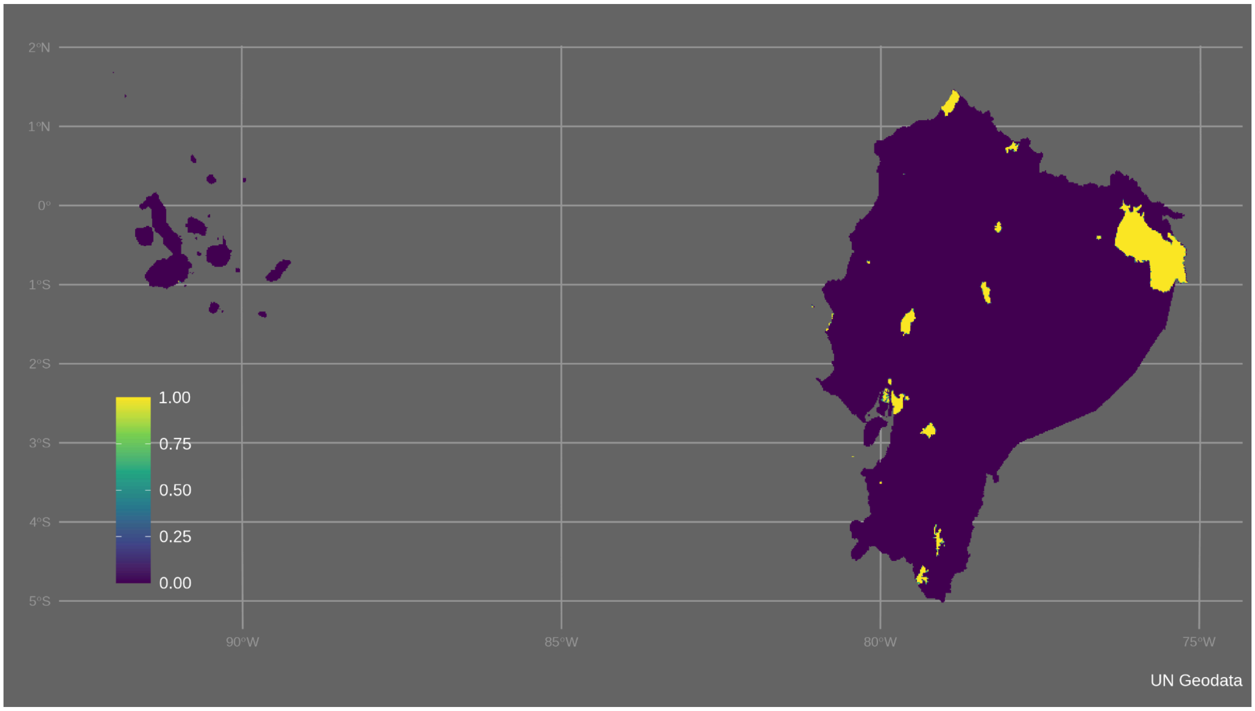

Figure S4. Planning features of ELSA Ecuador - RAMSAR Sites

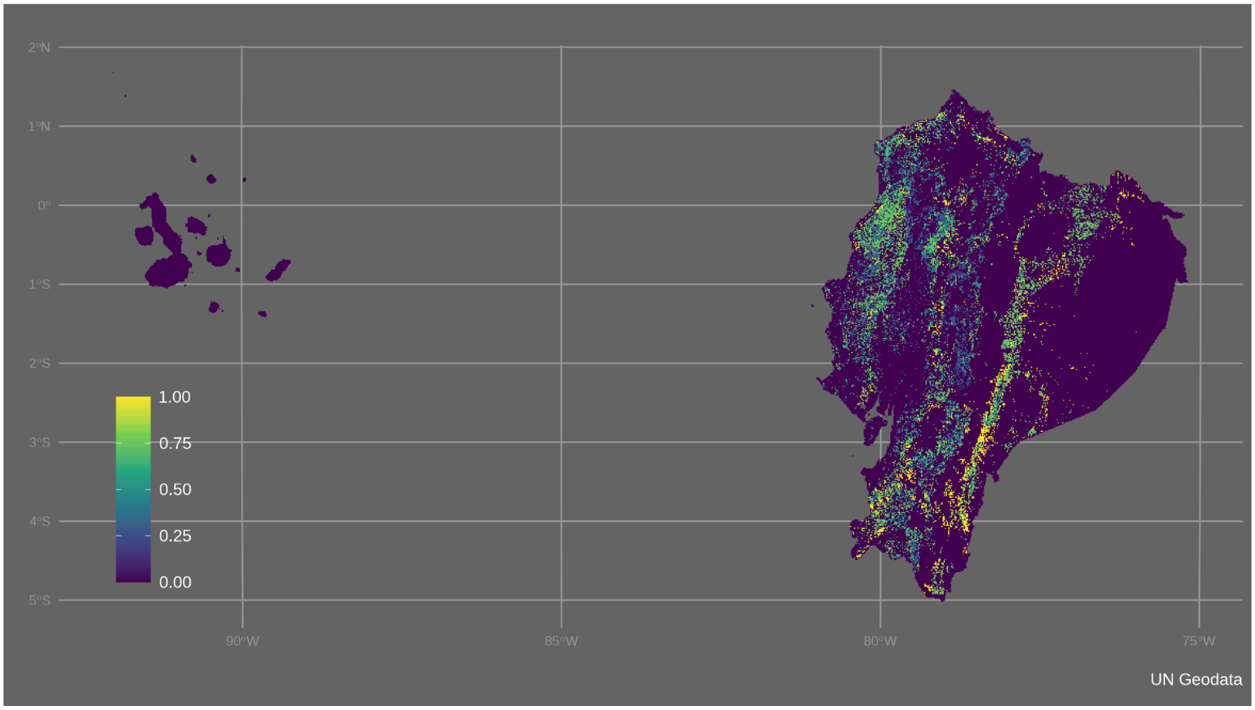


Figure S5. Planning features of ELSA Ecuador - forest restoration priority and agreement lands

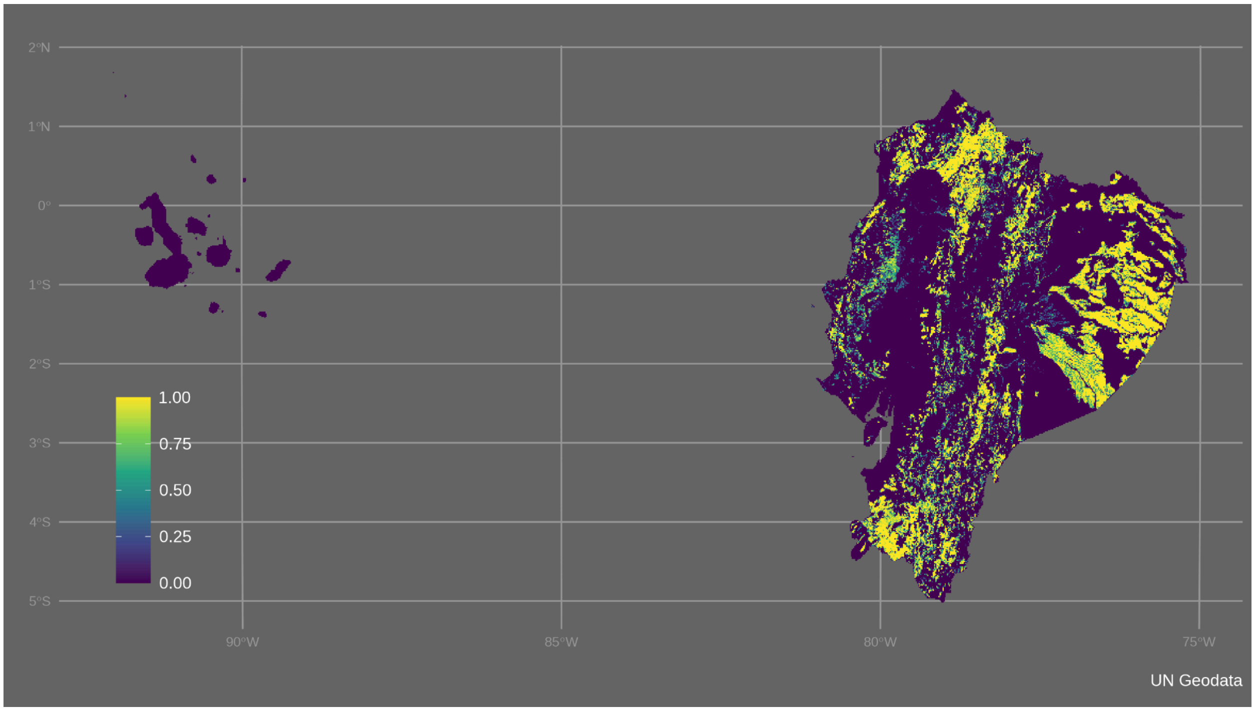


Figure S6. Planning features of ELSA Ecuador - forest for protection

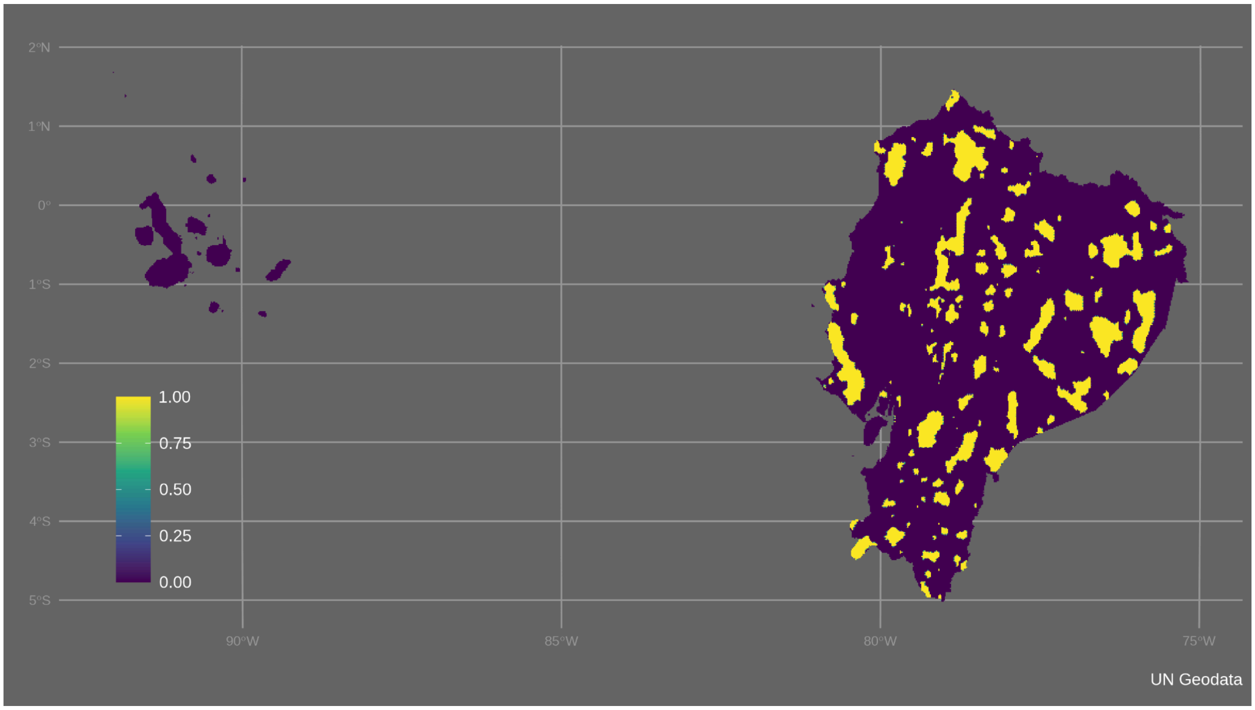


Figure S7. Planning features of ELSA Ecuador - priority biodiversity areas

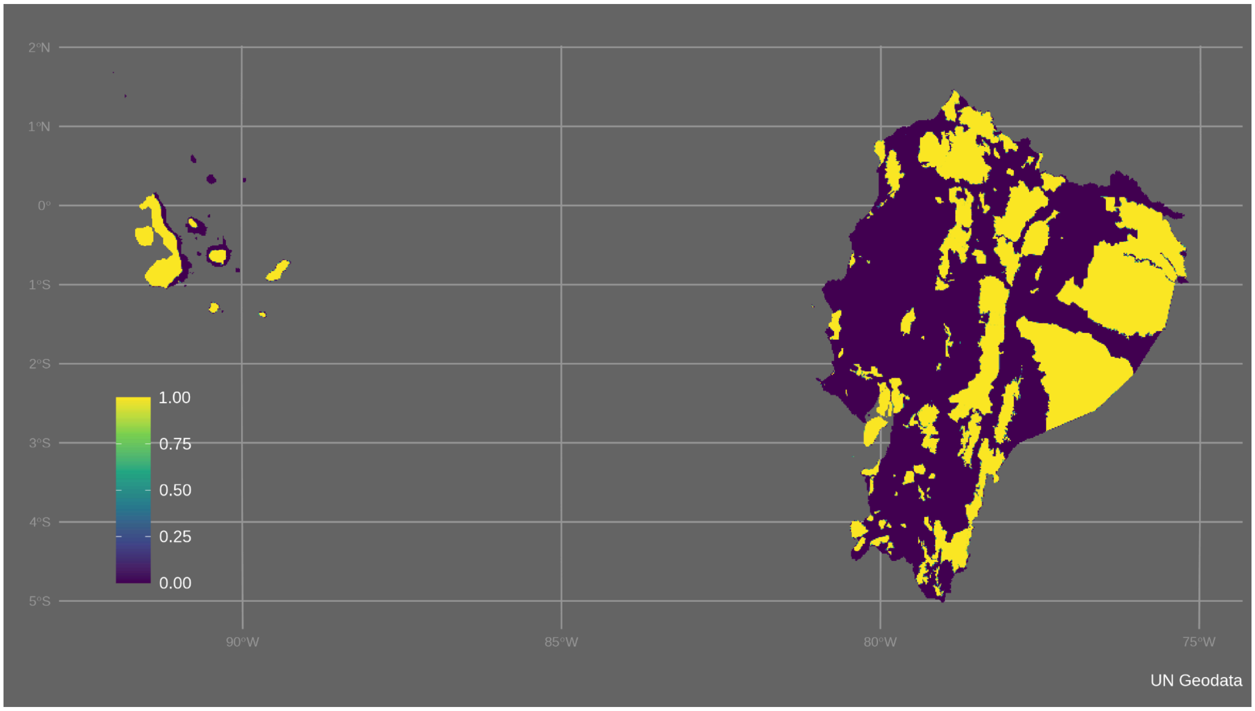


Figure S8. Planning features of ELSA Ecuador - KBAs

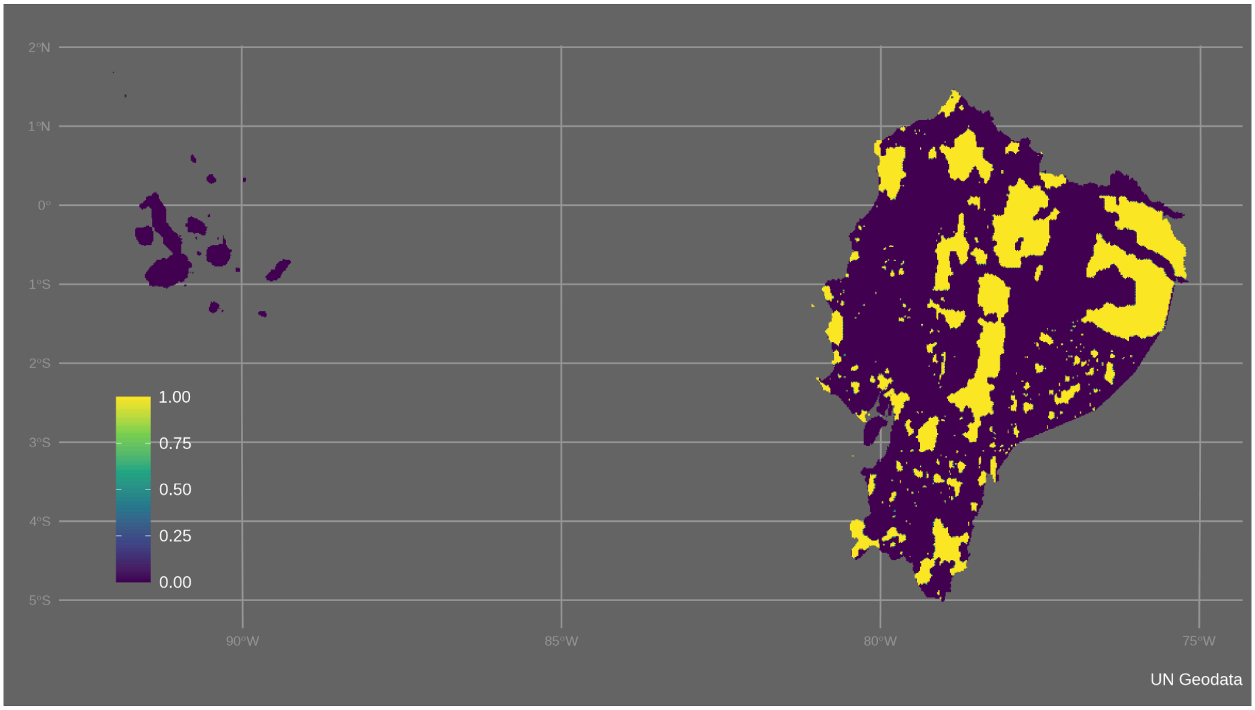


Figure S9. Planning features of ELSA Ecuador - conservation gaps

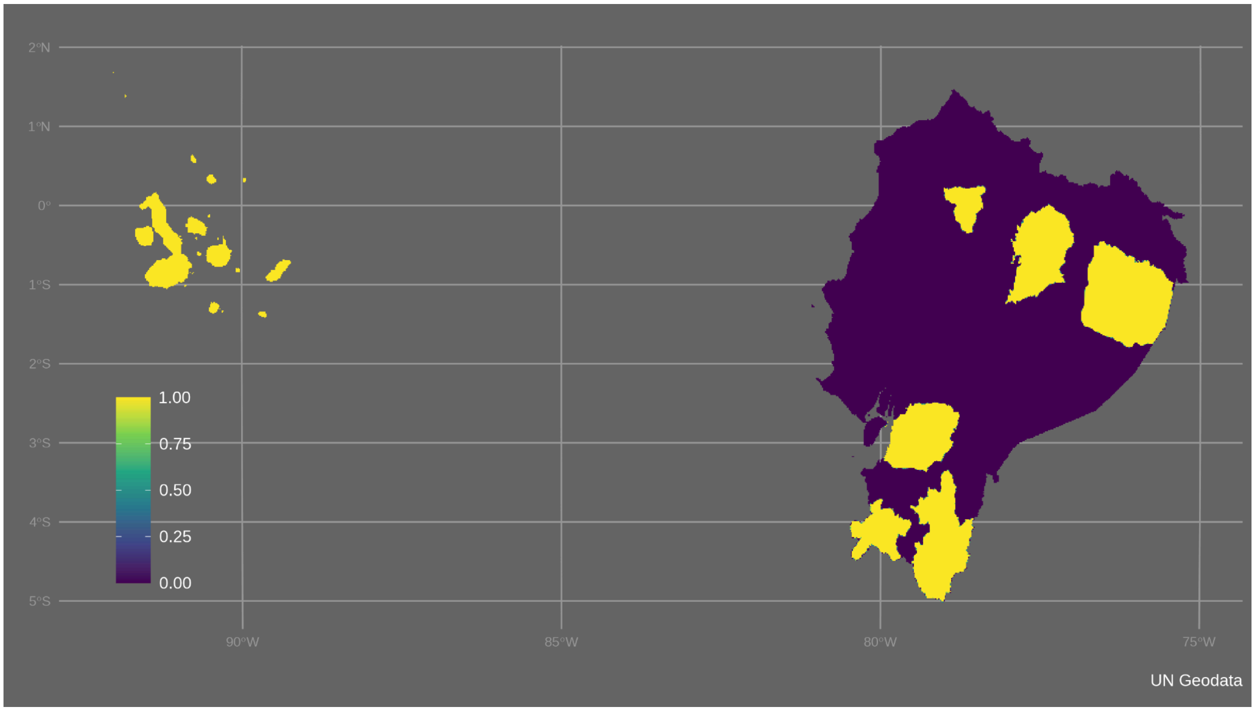


Figure S10. Planning features of ELSA Ecuador - biosphere reserves

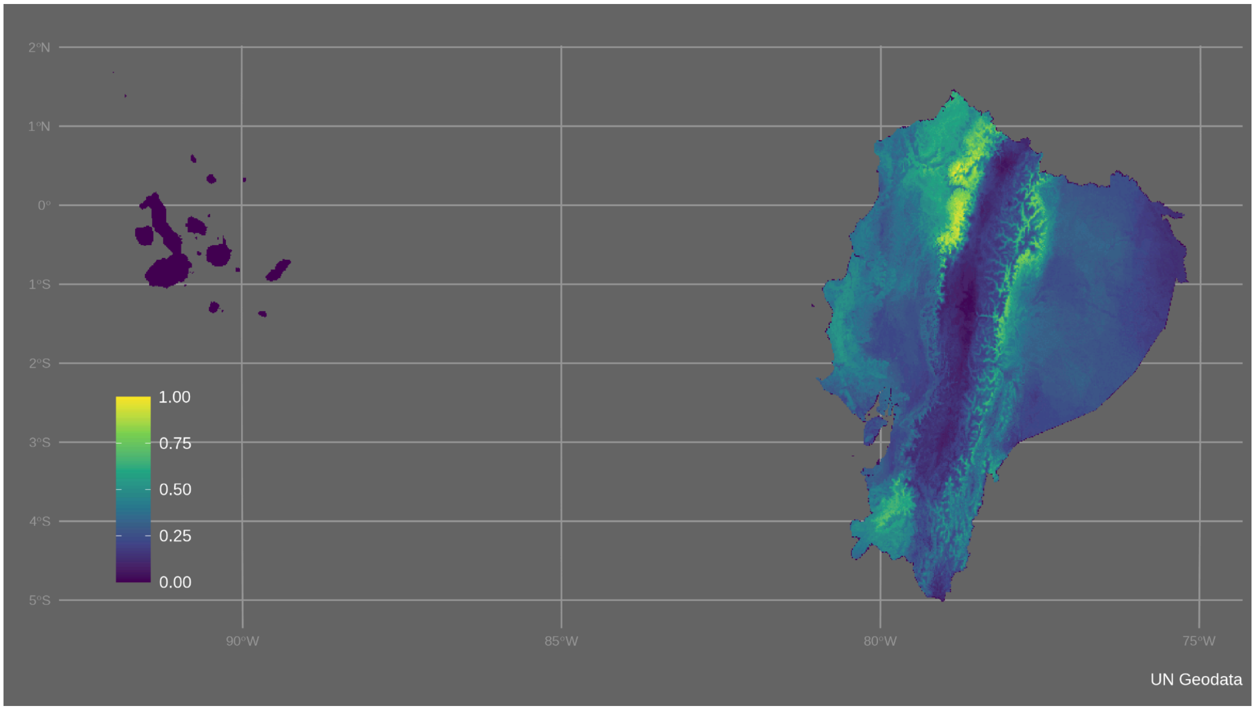


Figure S11. Planning features of ELSA Ecuador - bird species richness

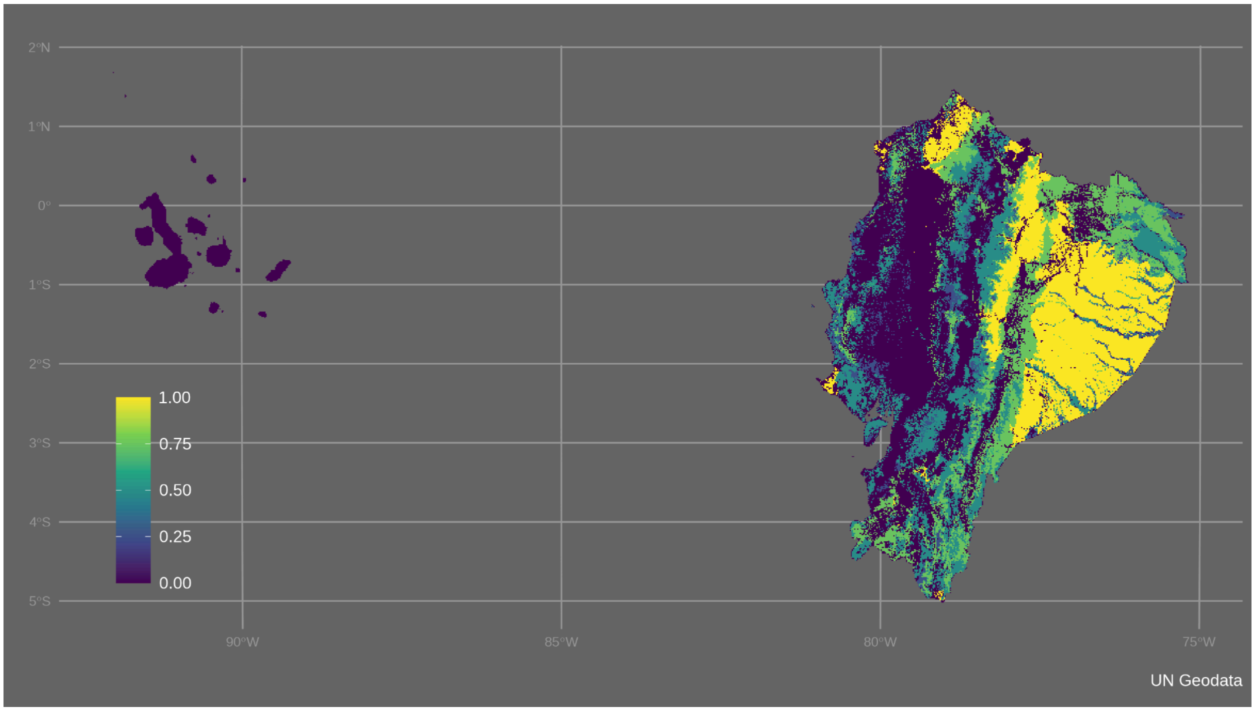


Figure S12. Planning features of ELSA Ecuador - connectivity of vegetation communities

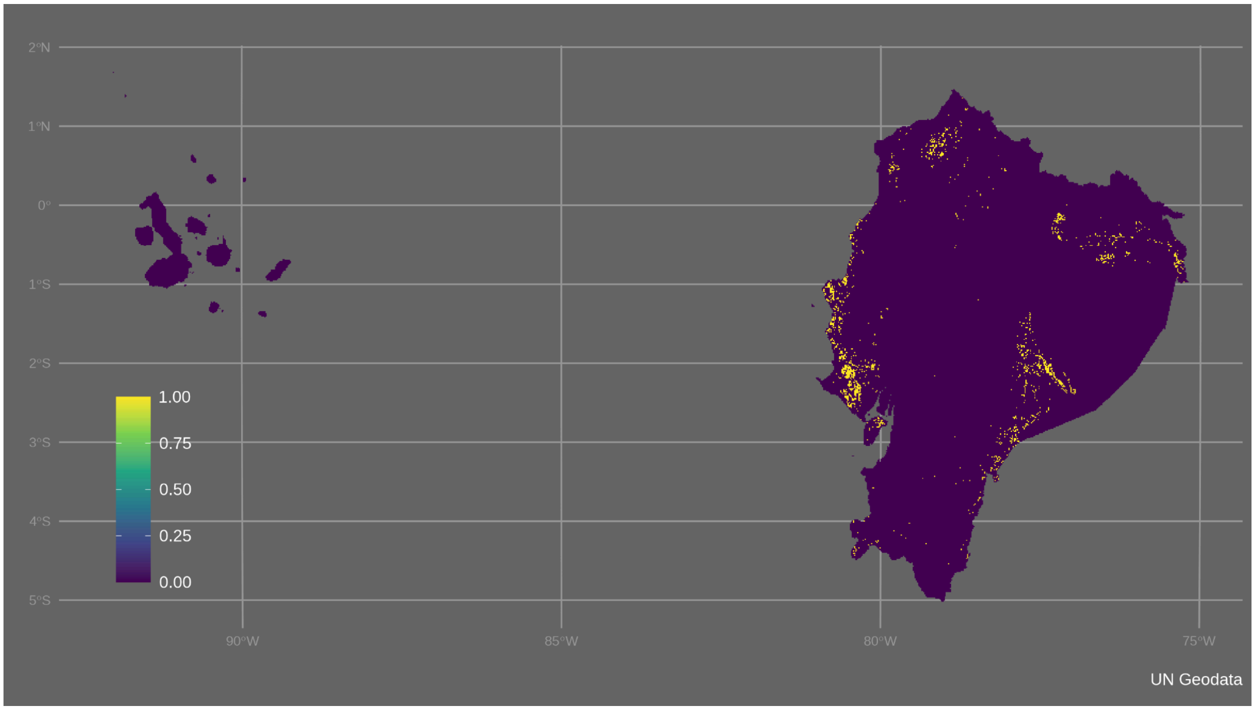


Figure S13. Planning features of ELSA Ecuador - vulnerable agricultural frontier

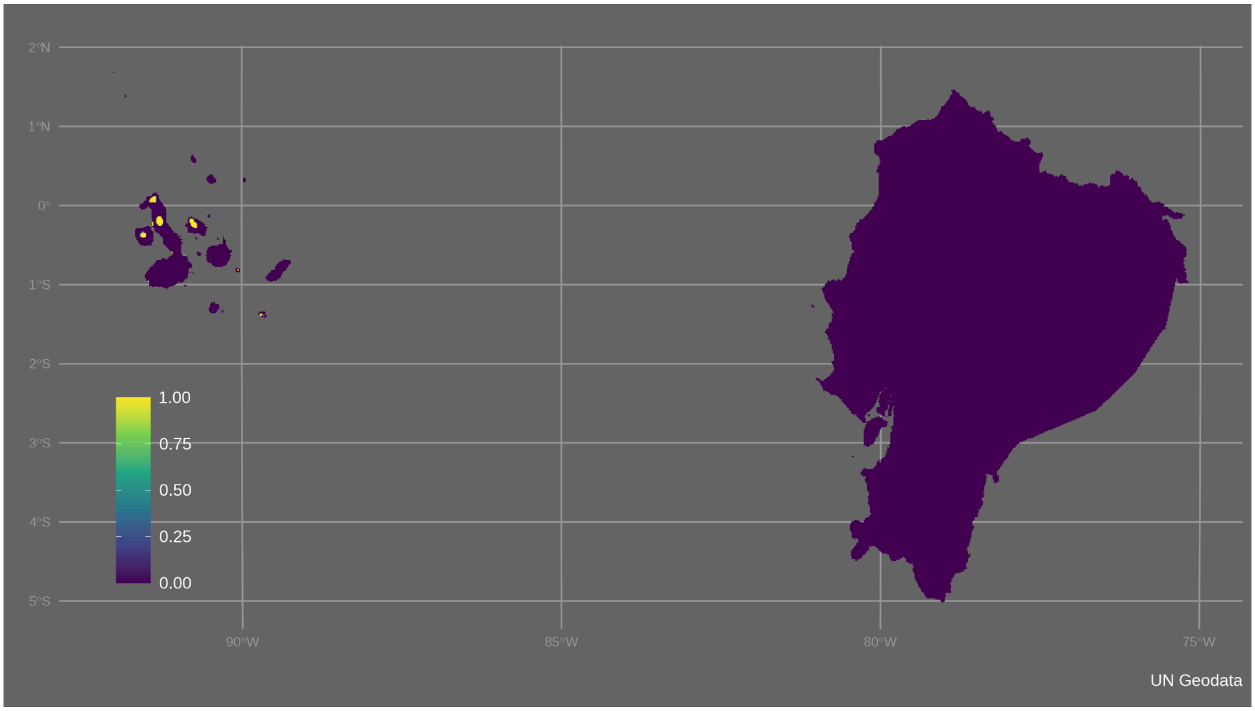


Figure S14. Planning features of ELSA Ecuador - Galapagos intangible zone

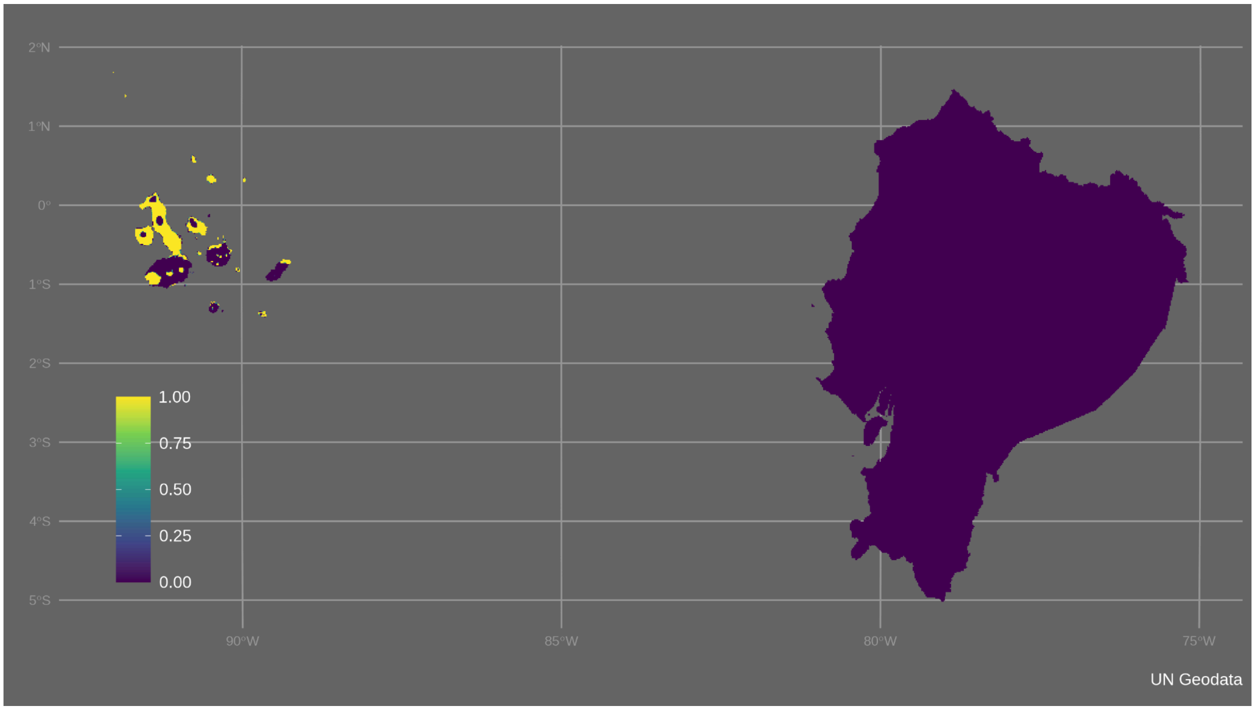


Figure S15. Planning features of ELSA Ecuador - Galapagos conservation zone

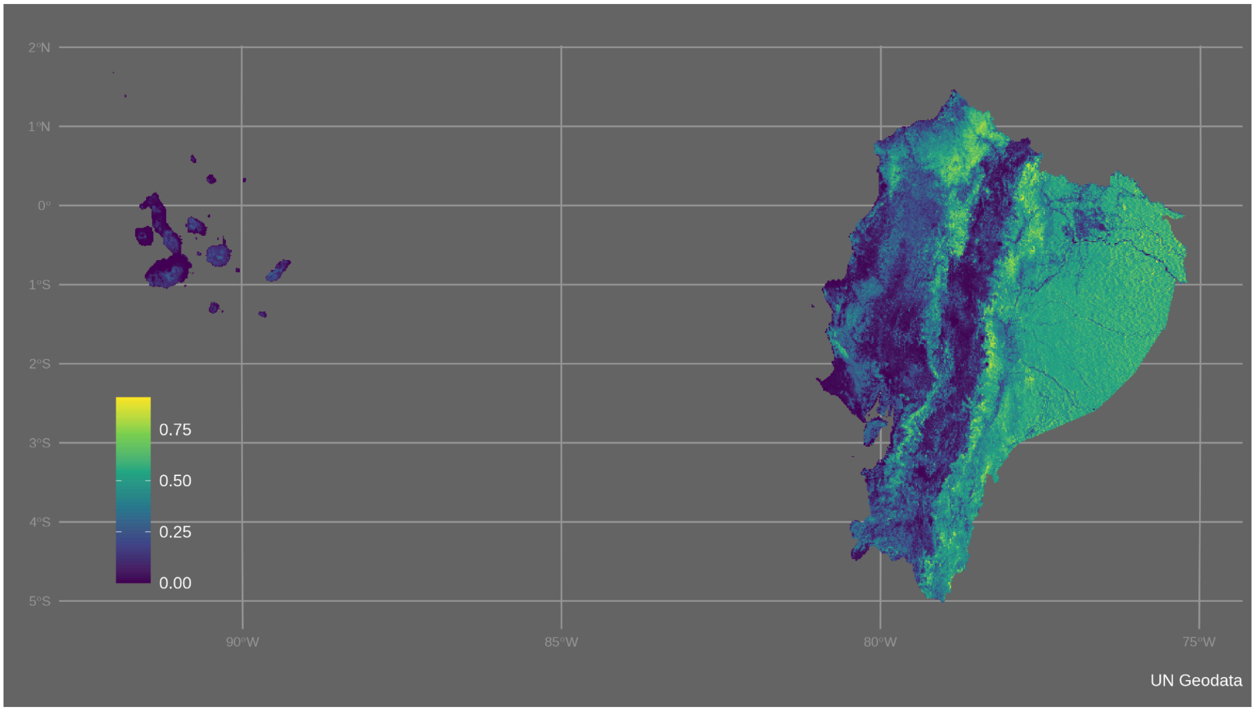


Figure S16. Planning features of ELSA Ecuador - biomass carbon

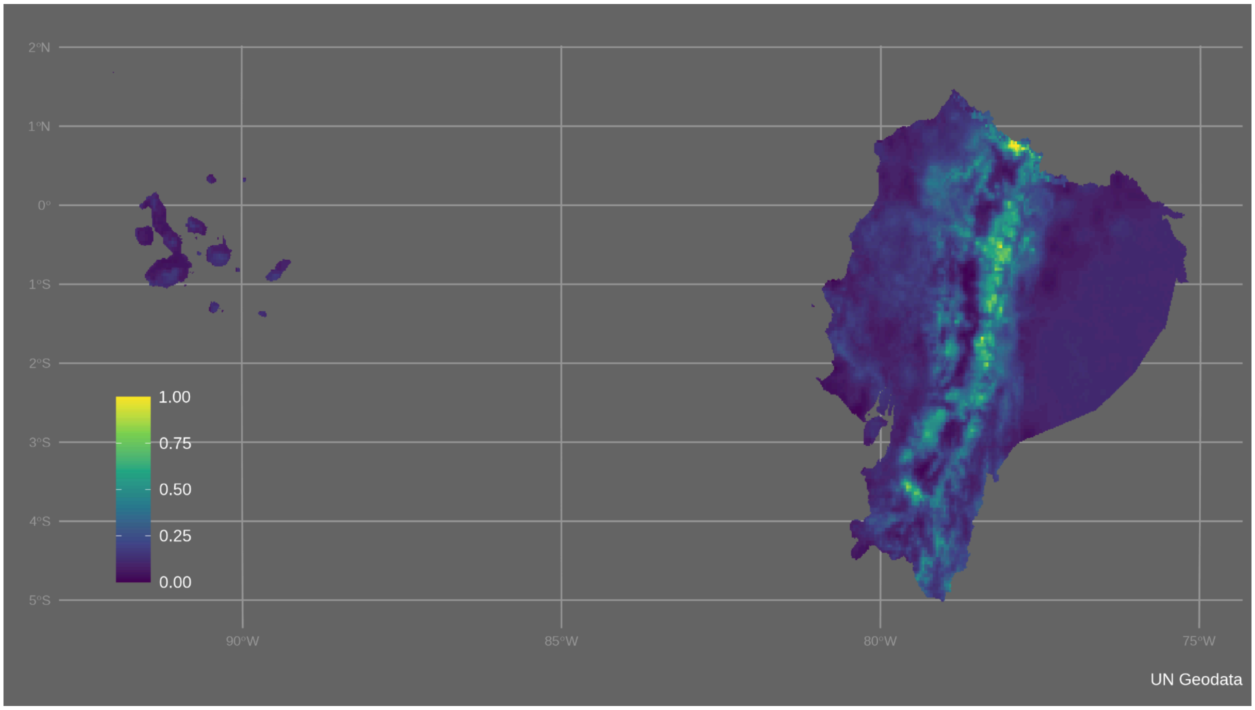


Figure S17. Planning features of ELSA Ecuador - soil organic carbon

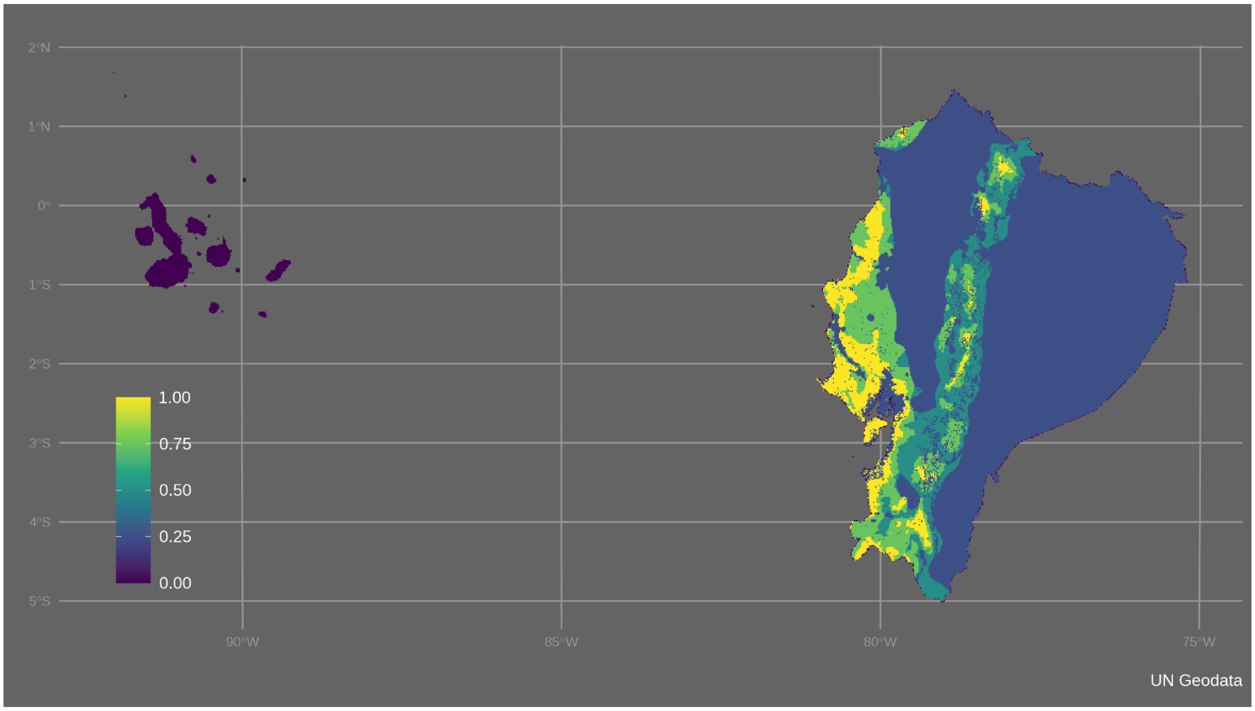


Figure S18. Planning features of ELSA Ecuador - susceptibility to desertification

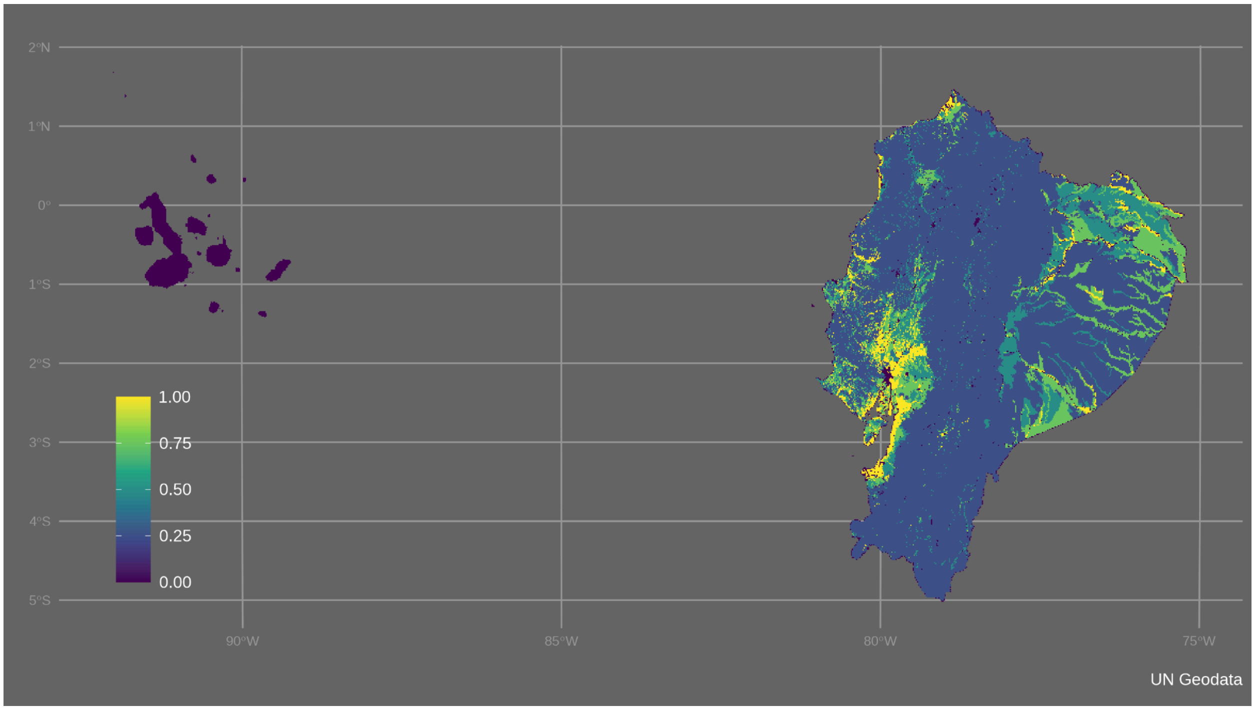

Figure S19. Planning features of ELSA Ecuador - susceptibility to flood

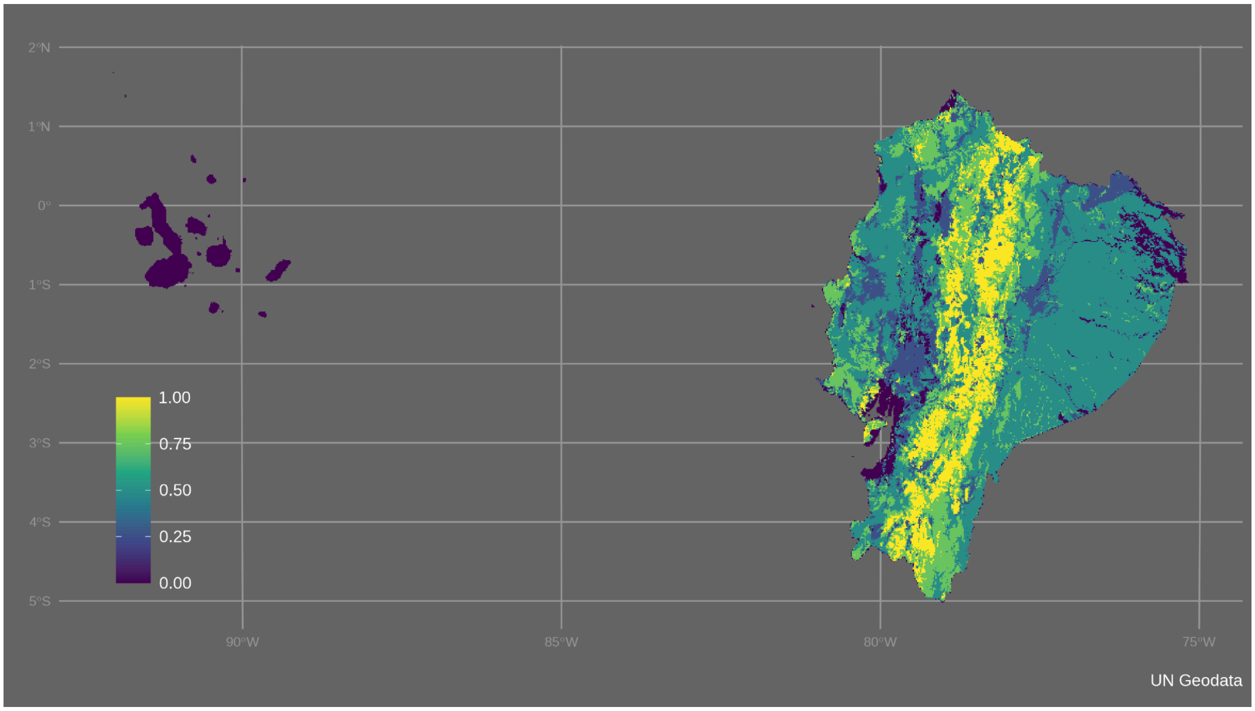

Figure S20. Planning features of ELSA Ecuador - susceptibility to forest fires

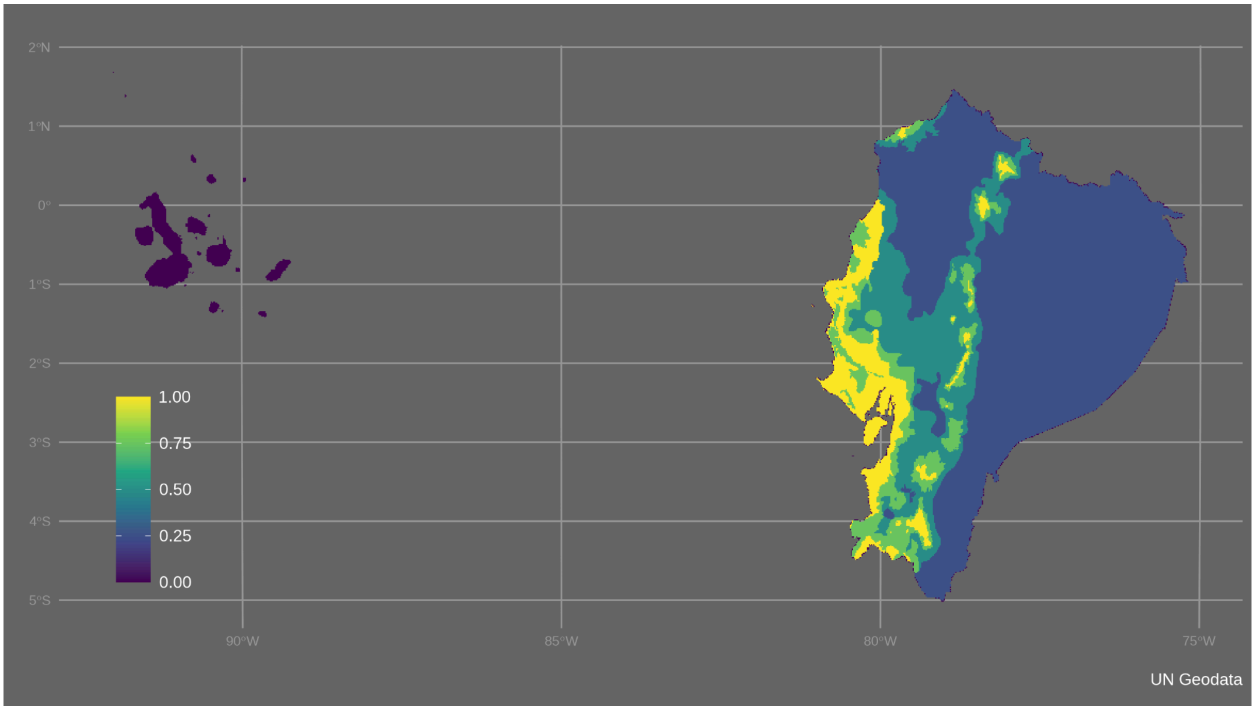

Figure S21. Planning features of ELSA Ecuador - susceptibility to drought

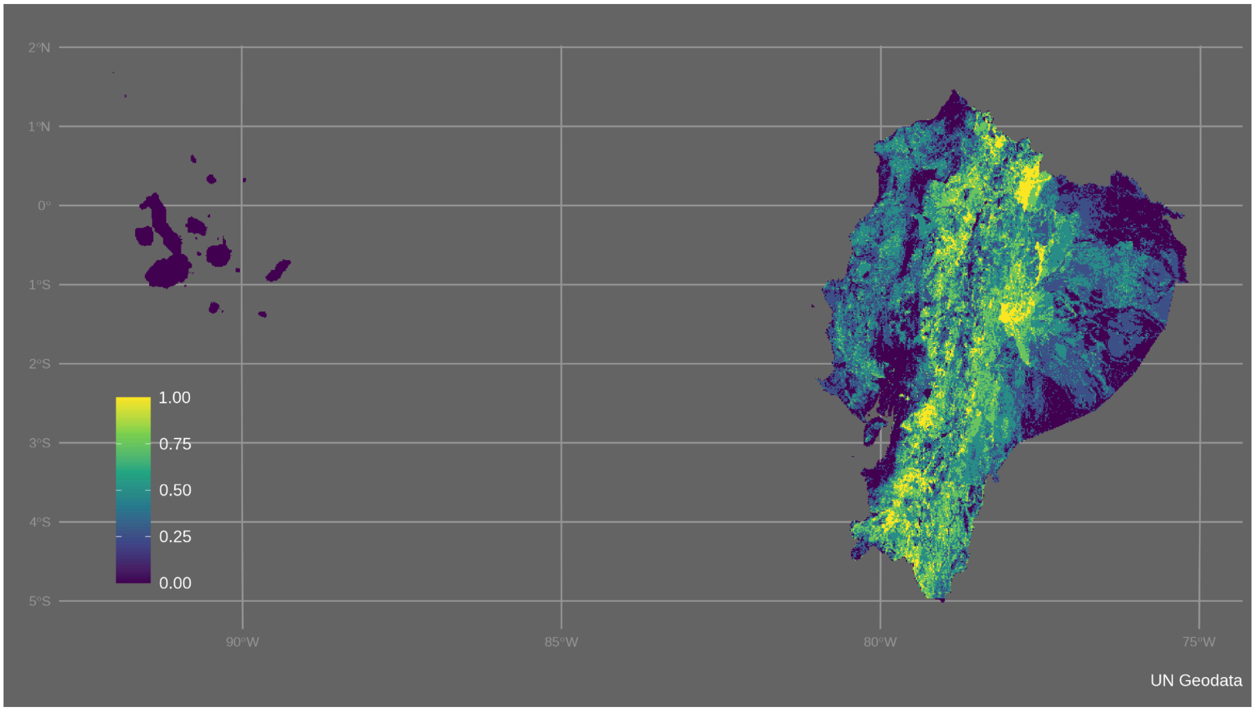

Figure S22. Planning features of ELSA Ecuador - susceptibility to mass movement

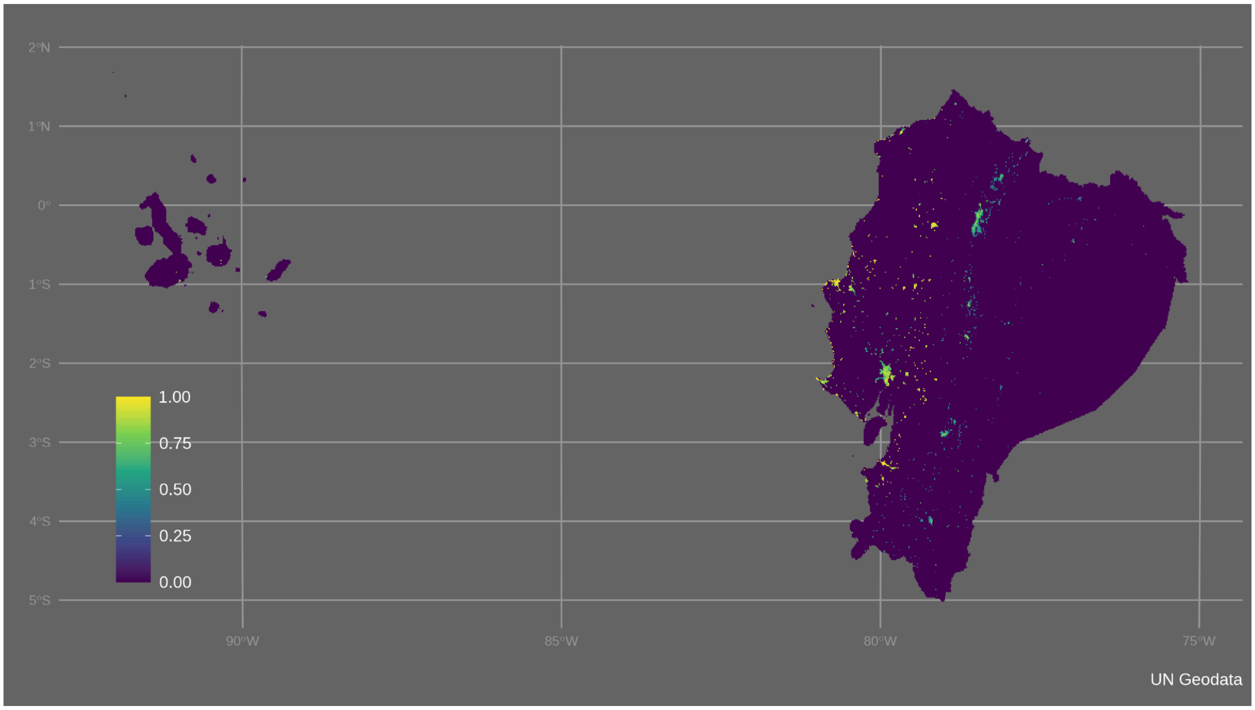


Figure S23. Planning features of ELSA Ecuador - urban-greening opportunities

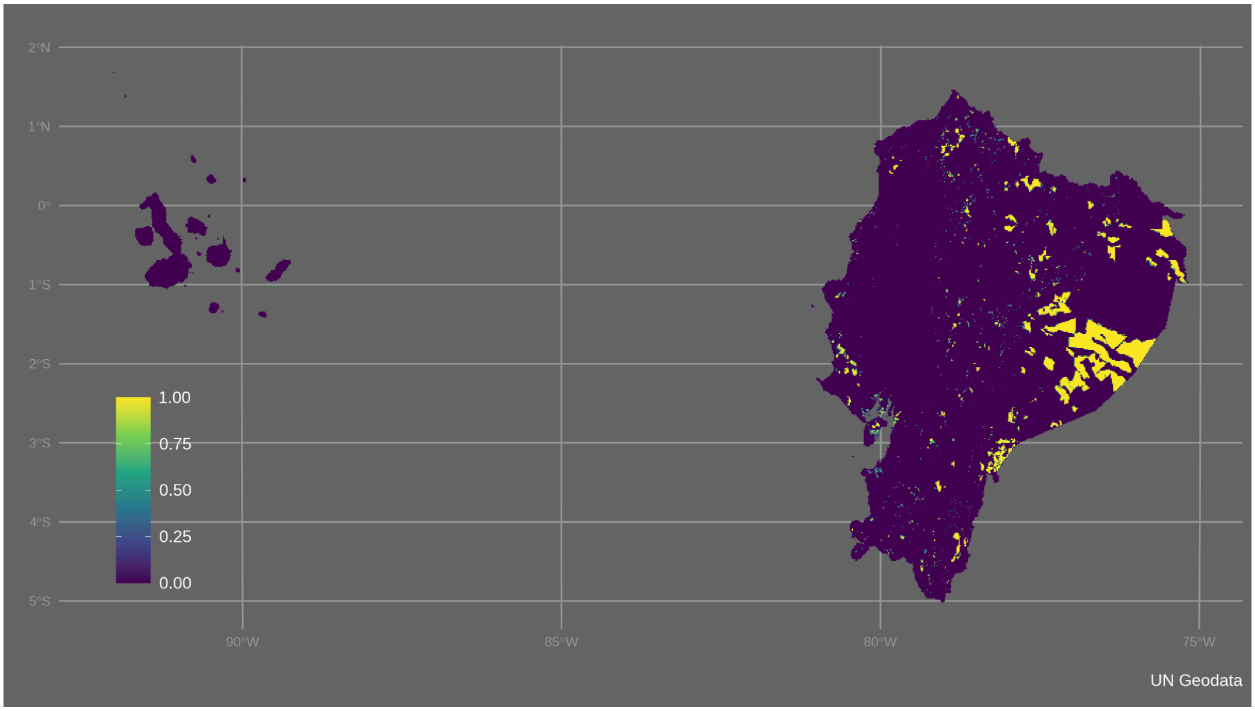


Figure S24. Planning features of ELSA Ecuador - socio-forest program area

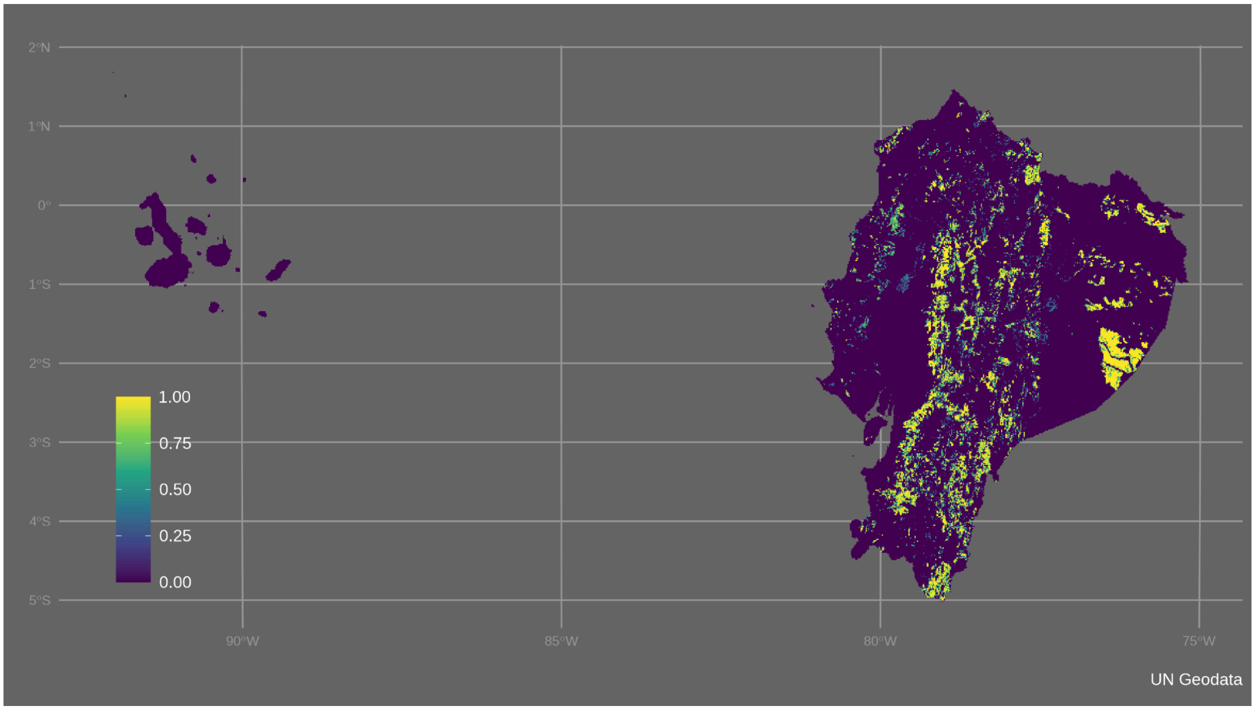

Figure S25. Planning features of ELSA Ecuador - forest for production

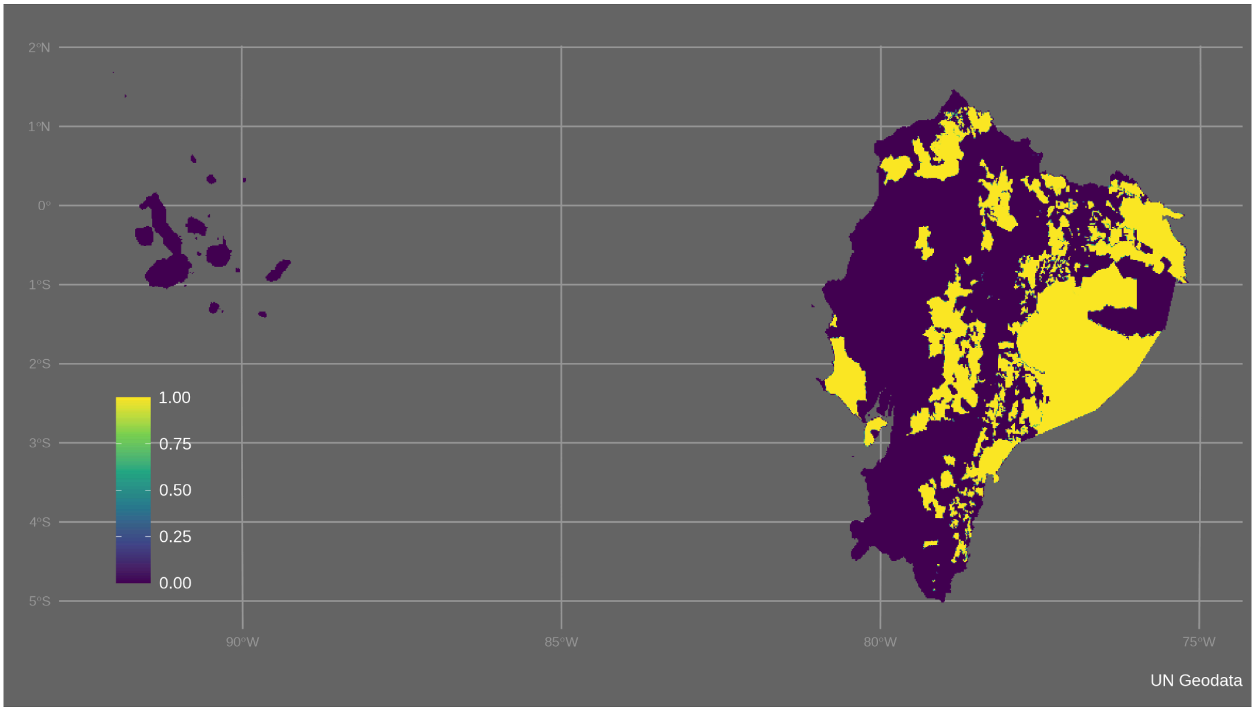

Figure S26. Planning features of ELSA Ecuador - indigenous territories

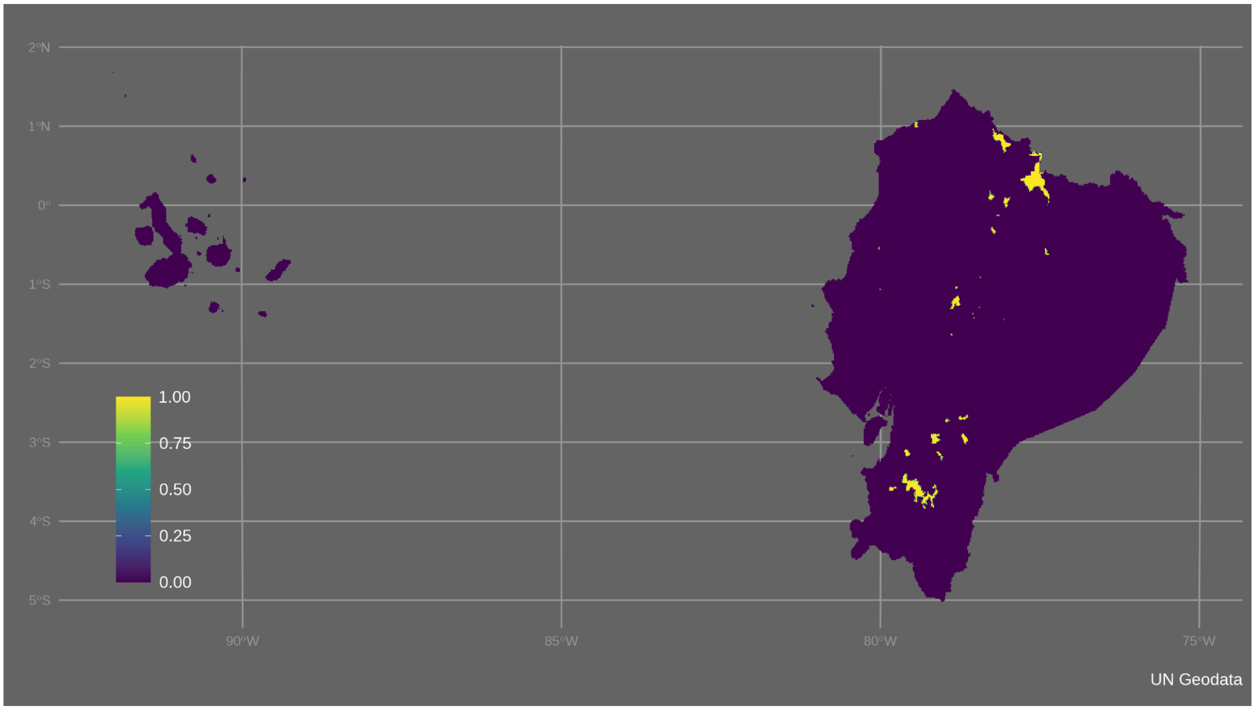

Figure S27. Planning features of ELSA Ecuador - important water source

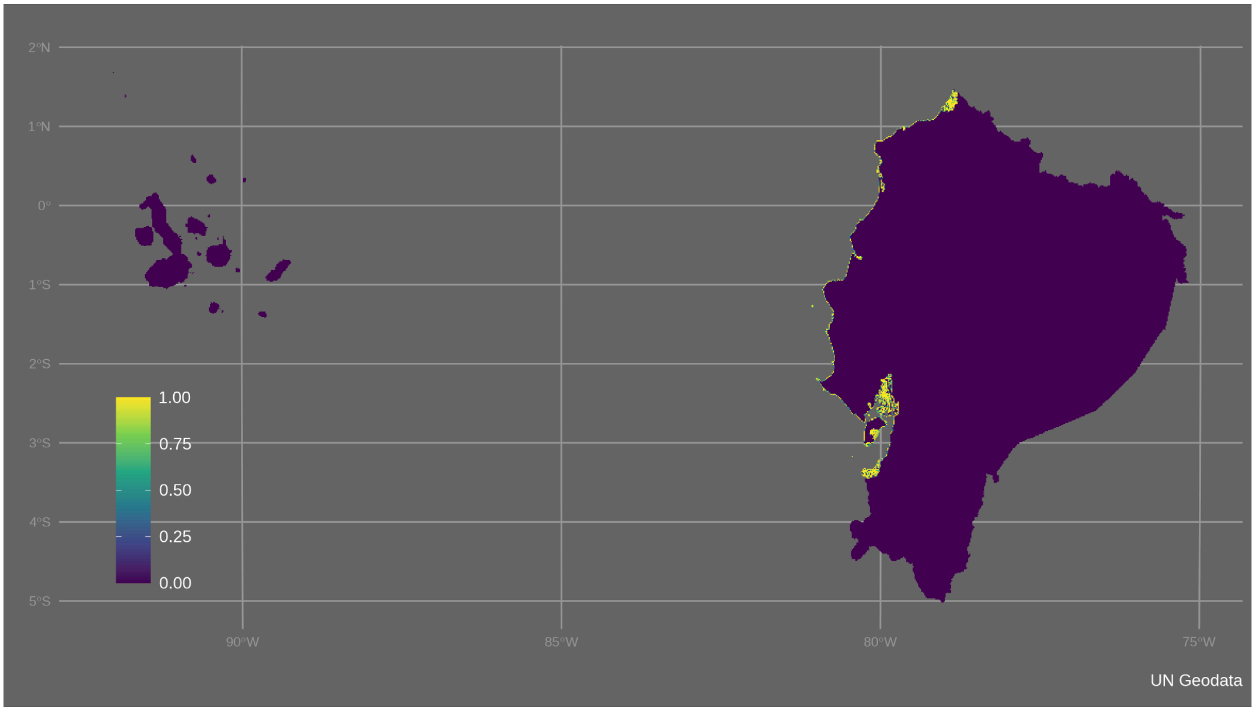

Figure S28. Planning features of ELSA Ecuador - coastal region

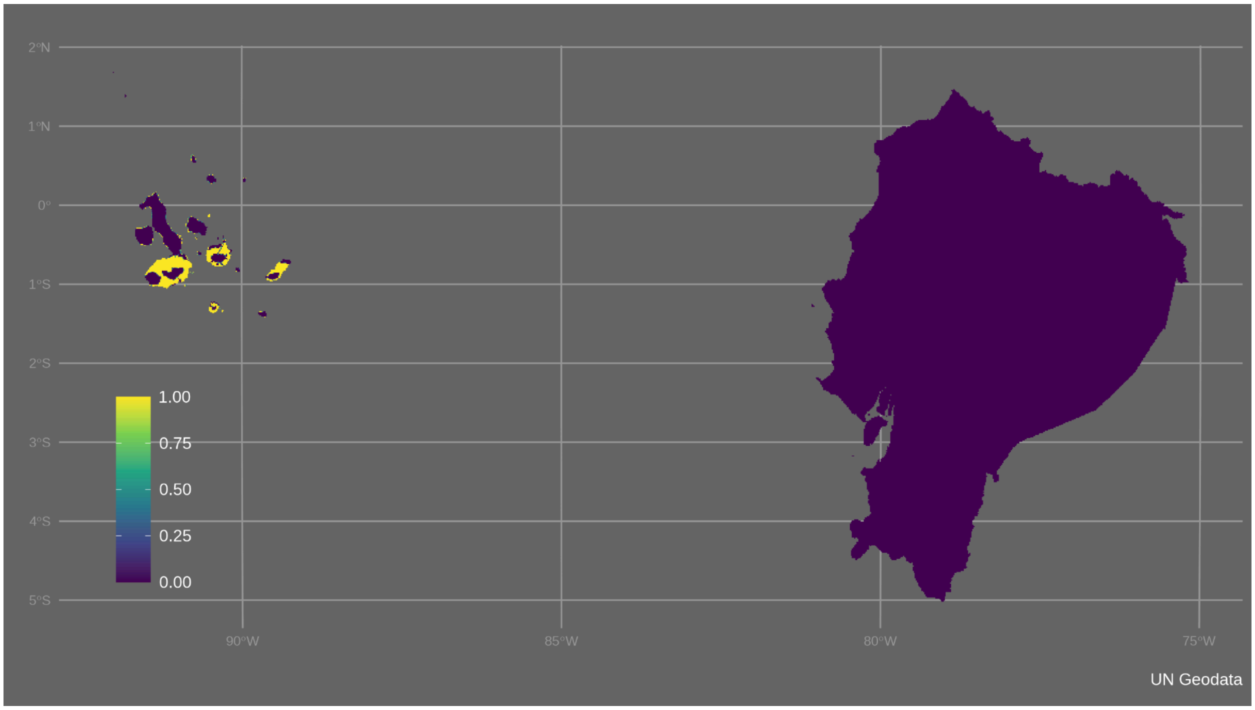


Figure S29. Planning features of ELSA Ecuador - Galapagos sustainable use

Table S2 Zone contribution scores of planning features in ELSA Ecuador

| Theme | Name | Protect Zone | Manage Zone | Restore Zone |
|---|---|---|---|---|
| Biodiversity | Native Forest | 1 | 0.5 | 1.5 |
| | Mangroves | 1 | 0 | 1.5 |
| | Paramos | 1 | 0.25 | 1.5 |
| | RAMSAR Sites | 1 | 0.25 | 1.5 |
| | Forest restoration priority and agreement lands | 0 | 0 | 1 |

| | | | | |
|---|---|---|---|---|
| | Forest for protection | 1 | 0 | 0 |
| | Priority biodiversity areas | 1 | 0.5 | 1.5 |
| | KBAs | 1 | 0.5 | 1.5 |
| | Conservation gaps | 1 | 0.5 | 1.5 |
| | Biosphere reserves | 1 | 1 | 1 |
| | Bird species richness | 1 | 0.5 | 1.5 |
| | Connectivity of vegetation communities | 1 | 0.5 | 0 |
| | Vulnerable agricultural frontier | 1 | 0.25 | 1.5 |
| | Galapagos intangible zone | 1 | 0 | 1 |
| | Galapagos conservation zone | 1 | 0 | 1 |
| Climate Change Mitigation | Biomass carbon | 1 | 0.5 | 1.5 |
| | Soil organic carbon | 1 | 0.75 | 1.5 |
| | Susceptibility to desertification | 1 | 0.5 | 1.5 |
| | Susceptibility to flood | 1 | 0.5 | 1.5 |
| | Susceptibility to forest fires | 1 | 0.5 | 1.5 |
| | Susceptibility to drought | 1 | 1 | 1.5 |

| | | | | |
|---|---|---|---|---|
| | Susceptibility to mass movement | 1 | 0.5 | 1.5 |
| | Urban greening opportunities | 0 | 0 | 1.5 |
| | Socio-forest program area | 1 | 1 | 1 |
| Sustainable Development | Forest for production | 0 | 1 | 0 |
| | Indigenous territories | 1 | 1 | 1.5 |
| | Important water source | 1 | 0.5 | 1.5 |
| | Coastal region | 1 | 0.5 | 1.5 |
| | Galapagos sustainable use zone | 0 | 1 | 0 |

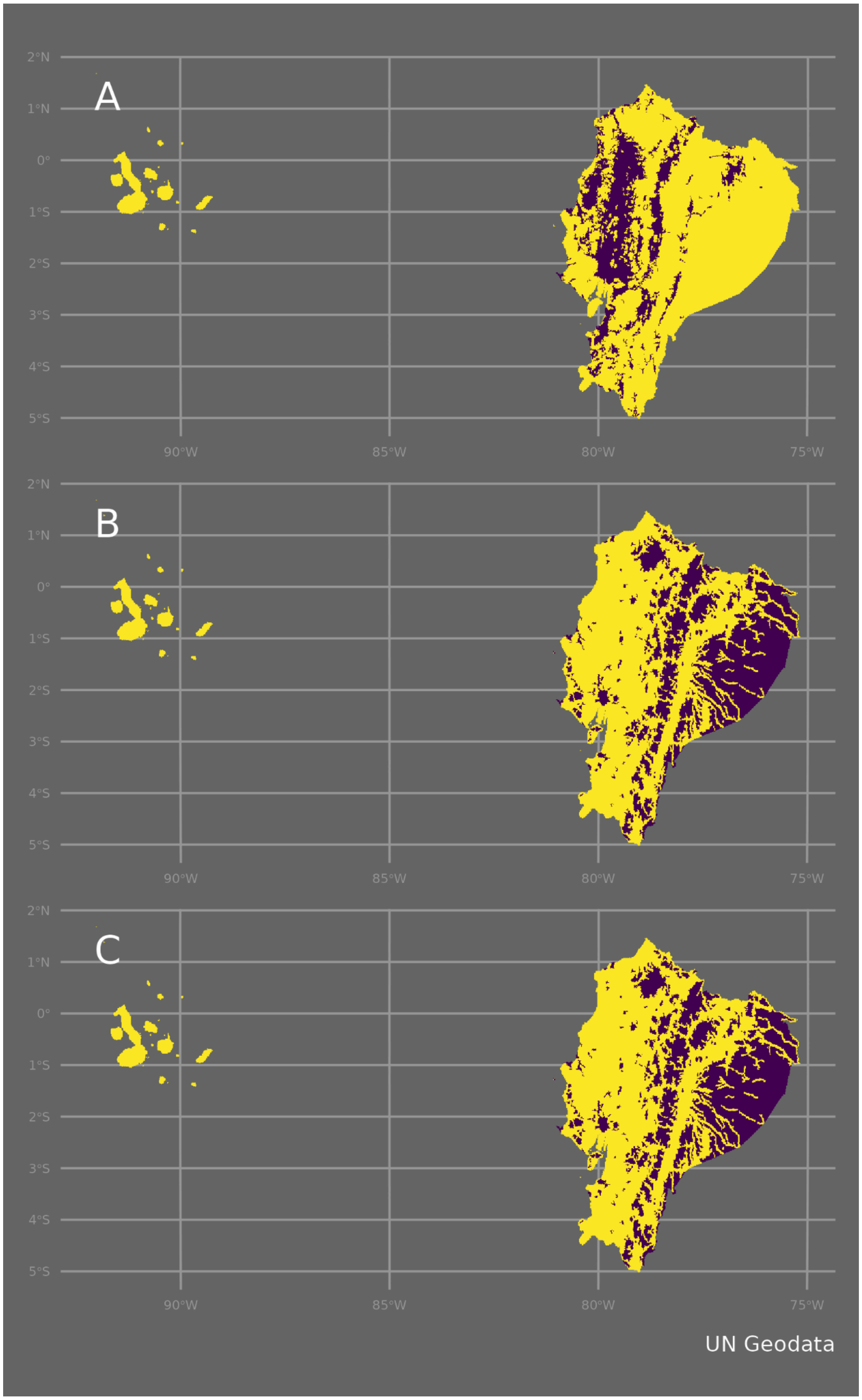


Figure S30. Nature-based action zones of ELSA Ecuador, indicating the extent of the (A) protection zone, (B) restoration zone, and (C) management zone (areas are in yellow). Restoration and management zones share the same spatial restrictions in ELSA Ecuador.

Table S3. Spatial constraints of nature-based action zones in ELSA Ecuador

| Zones | Spatial constraints | Coverage of planning areas |
|---|---|---|
| Protect | Human Footprint Index <6, representing 95% of the index's distribution within existing PAs. | 80.04% |
| Manage | Human Footprint Index > 0 and <10, representing the middle 60% of the index's distribution outside of protected areas in the country | 67.27% |
| Restore | Human Footprint Index > 0 and <10, representing the middle 61% of the index's distribution outside of protected areas in the country | 67.27% |
